# Novel Small Molecules Targeting the Intrinsically Disordered Structural Ensemble of α-Synuclein Protect Against Diverse α-Synuclein Mediated Dysfunctions

**DOI:** 10.1101/646505

**Authors:** Gergely Tóth, Thomas Neumann, Amandine Berthet, Eliezer Masliah, Brian Spencer, Jiahui Tao, Michael F. Jobling, Shyra J. Gardai, Carlos W. Bertoncini, Nunilo Cremades, Michael Bova, Stephen Ballaron, Xiao-Hua Chen, Wenxian Mao, Phuong Nguyen, Mariano C. Tabios, Mitali A. Tambe, Jean-Christophe Rochet, Hans-Dieter Junker, Daniel Schwizer, Renate Sekul, Inge Ott, John P. Anderson, Balazs Szoke, Wherly Hoffman, John Christodoulou, Ted Yednock, David A. Agard, Dale Schenk, Lisa McConlogue

## Abstract

The over-expression and aggregation of α-synuclein (αSyn) are linked to the onset and pathology of Parkinson’s disease. Native monomeric αSyn exists in an intrinsically disordered ensemble of interconverting conformations, which has made its therapeutic targeting by small molecules highly challenging. Nonetheless, here we successfully target the monomeric structural ensemble of αSyn and thereby identify novel drug-like small molecules that impact multiple pathogenic processes. Using a surface plasmon resonance high-throughput screen, in which monomeric αSyn is incubated with microchips arrayed with tethered compounds, we identified novel αSyn interacting drug-like compounds. Because these small molecules could impact a variety of αSyn forms present in the ensemble, we tested representative hits for impact on multiple αSyn malfunctions *in vitro* and in cells including aggregation and perturbation of vesicular dynamics. We thereby identified a compound that inhibits αSyn misfolding and is neuroprotective, multiple compounds that restore phagocytosis impaired by αSyn overexpression, and a compound blocking cellular transmission of αSyn. Our studies demonstrate that drug-like small molecules that interact with native αSyn can impact a variety of its pathological processes. Thus, targeting the intrinsically disordered ensemble of αSyn offers a unique approach to the development of small molecule research tools and therapeutics for Parkinson’s disease.

## Introduction

The sequential misfolding of α-synuclein (αSyn) into oligomers and fibrils is central to the pathogenesis of Parkinson’s Disease (PD) and related neurodegenerative disorders termed synucleinopathies^1^. Lewy bodies containing αSyn amyloid-like fibril inclusions are a hallmark neuropathological feature of these diseases^2^. The severity of disease correlates with the progressive spread of aggregated αSyn in patients^3^, and αSyn misfolding is associated with toxicity in cell and animal models^4^. In addition, strong genetic evidence links αSyn to PD including gene multiplications or missense mutations that cause rare early onset forms of PD^5–7^ and genetic association studies also link αSyn to sporadic PD^8,9^. This combined neuropathological, biochemical and genetic evidence provides strong support implicating the misfolding and aggregation of αSyn as a key feature in PD.

Monomeric αSyn, as an intrinsically disordered protein (IDP), exists primarily as a dynamic ensemble of distinct interconverting conformations that have the ability to take on more structured forms under the appropriate cellular context^10,11^. In particular, αSyn can take on more ordered forms upon membrane binding^12–14^. Owing to this intrinsic dynamic character and the ability to populate other forms upon interaction, IDPs are involved in many key biological processes^15^, are vulnerable to aggregation and are susceptible to prion-like amplification of misfolded species and transmission between cells^16^. This propensity towards aggregation and spread is likely to underlie the association that IDPs have with a growing number of misfolding diseases, notably many neurodegenerative disorders^17^.

Small molecule binding to native states of globular proteins has been successfully used to block misfolding and aggregation^18^ most notably in the case of targeting transthyretin to treat systemic amyloidosis^19^. By contrast, targeting of IDPs such as native monomeric αSyn with small molecules has been challenging due to their inherent structural heterogeneity and the absence of persistent structural elements^10^. Yet, the dynamic nature of the monomeric form of αSyn also provides opportunity to impact multiple aspects of the protein. For example, small molecules interacting with native αSyn could protect and stabilize specific conformations present in the ensemble, which in turn could enhance or inhibit misfolding, or modulate cellular malfunction associated with αSyn overexpression.

In spite of the inherent challenges, we previously identified potential small molecule binders to αSyn using an *in silico* screen. We showed that one of these compounds, 484228, displayed protective activity in reversing αSyn overexpression mediated neurodegeneration and impairment of phagocytosis without impact on aggregation of the recombinant protein^20^, thus supporting the notion that small molecules can target an IDP such as αSyn and in doing so modulate cellular malfunctions independent of inhibitory effects on aggregation in solution. Encouraged by the success of the *in silico* screen, we embarked on biophysical screens to identify compounds that bind to IDPs using a surface plasmon resonance (SPR) based assay in which compounds are tethered to the chip (high-throughput chemical microarray surface plasmon resonance imaging, HT-CM-SPR)^21^. HT-CM-SPR has been successfully applied for the identification of small molecule binders to globular proteins, providing starting hits for drug discovery^21,22^. Remarkably, we successfully used HT-CM-SPR to identify small molecules retarding the aggregation of tau protein, another IDP that misfolds in neurodegenerative diseases^23^.

Here we apply the HT-CM-SPR screening technology to αSyn, and in addition to searching for aggregation-blocking compounds, as we did in the tau screen, we search for compounds correcting additional disease-relevant malfunctions of αSyn. We report here the identification of novel compounds that interact with native αSyn. A subset of these compounds can rescue αSyn dysfunction by reducing αSyn aggregation. Others can restore vesicular dynamics impaired by αSyn overexpression, as reflected in phagocytic capacity, while a distinct compound can block αSyn cell-to-cell transmission, without direct impact on aggregation. The identification of small molecules reversing diverse malfunctions of αSyn indicates that differing conformations and associated malfunctions of the protein may be targeted by small molecule ligands.

## Results

### Identification of small molecule binders of αSyn by high-throughput chemical microarray surface plasmon resonance imaging (HT-CM-SPR) screening

Monomeric αSyn was screened against a library of small molecules containing 91,000 lead-like and 23,000 fragment compounds immobilized on microarrays to identify small molecules binding to the protein using surface plasmon resonance (SPR) imaging (HT-CM-SPR)^21,22^ (Fig. 1a). An advantage of this chemical microarray paradigm is that αSyn is maintained in a soluble, monomeric, and label-free state enabling it to assume its heterogeneous conformational ensemble during screening. Moreover, the SPR-based detection is highly sensitive, which allows for identifying weak binding events^22^.

**Figure 1.**
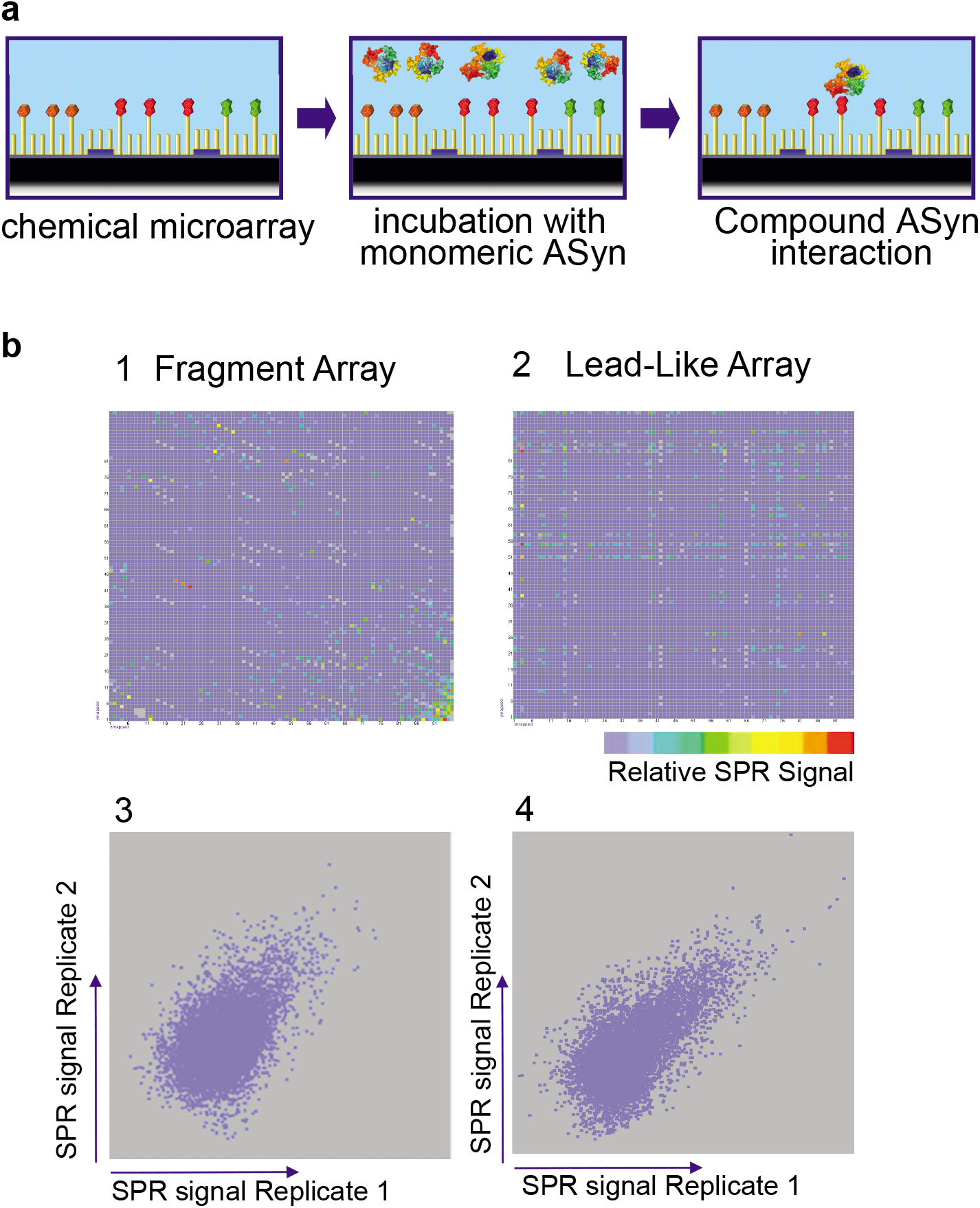
HT-CM-SPR Screening of αSyn. **a)** The HT-CM-SPR process: Monomeric αSyn analyte floats over the array surface in screening buffer to allow binding events to occur. SPR imaging enables the detection of binding events. **b).** SPR Test Screening Results: The upper two panels show images of color coded SPR signals obtained during HT-CM-SPR of αSyn under optimized screening conditions against individual representative microarrays comprised of 1) a fragment microarray containing 3,070 fragments (out of 23,000 fragments present in the entire screened NovAliX chemical microarray library) spotted in triplicate along a diagonal and 2) a lead-like microarray of 9,216 lead-like compounds (out of 91,000 lead-like compounds present in the entire screened NovAliX chemical microarray library) individually spotted. The lead-like compounds are derived from combinatorial synthesis approaches with each row and column on the microarrays consisting of a common fragment in combination with 96 other diversities. In these fingerprint representations of the microarrays each spot is representative of a separate small molecule tethered to the array with the location on the chip reflected by the location on the fingerprint and the color representing the intensity of signal for that spot (color ranging from low shifts (blue) to high (red) signals). The lower panels 3 and 4 show reproducibility of duplicate screening experiments for chemical microarrays shown in panels 1 and 2. The reproducibility of these subsets of compounds is representative of results obtained for the entire NovAliX chemical microarray library with 3) triplicates of 3,070 fragments and for 4) individual 9,216 lead-like compounds immobilized. In these scatter plots, each spot compares the signal strength measured for the microarray samples in each of the independent array experiments with signals from one replicate experiment indicated on the Y-axis and that of the other experiment represented on the X-axis.

The monomeric nature of the αSyn was verified by dynamic light scattering (DLS), its native monomeric conformation ensemble by Nuclear Magnetic Resonance (NMR) and its purity by electrophoretic analyses (Supplementary Figs. S1-S3). NMR H^1^-N^15^-HSQC analyses compared the commercially generated αSyn protein used in the screen to that prepared in-house using standard protocols for preparation of monomeric ensembles of αSyn and they showed identical spectra (Supplementary Fig. S2). To verify the monomeric nature of αSyn in the screen, DLS analyses were performed on αSyn preparations under screening conditions and in parallel for each library screening experiment. αSyn stock solutions were shown to be free of aggregates with negligible (<0.1%,) amounts of oligomers present. αSyn in screening buffer subjected to three-hour long incubations, the maximum amount of time used during screening, remained monomeric (Supplementary Fig. S3). In addition, there was no evidence of increasing SPR signal during the three-hour screening, as would have been seen had the αSyn been oligomerizing on the microchip. Thus, the detected SPR signal reflects the interactions between monomeric αSyn and the immobilized library.

Microarrays were incubated with αSyn under optimized conditions, imaged and subjected to hit selection as described in the Supplementary Information. Briefly, chips were incubated with monomeric αSyn under conditions optimizing signal and shown to maintain αSyn in its monomeric state (Supplementary Fig. S3). Fig 1b shows representative examples of αSyn SPR fingerprint analyses for two individual microarrays selected from the 13 arrays that comprise the library. Shown are examples of a microarray containing fragments (panels 1 and 3) and another microarray containing lead-like compounds (panels 2 and 4). Panels 1 and 2 show imaging analyses of the microarrays with each spot reflecting the position of a tethered small molecule sample on the array and the color of that spot reflecting the signal upon incubation with αSyn. On chips containing leadlike compounds based on combinatorial synthesis approaches, related library compounds are arrayed in rows and columns, with one unique copy per compound species present per physical microarray. Hence hit series with multiple members interacting with αSyn generated array patterns of colored lines as shown in Fig. 1b panel 2. Such patterns can be used to identify hit series with multiple related compounds interacting with the target protein. Some chips also contain many singleton compounds for diversity in the library, so these generated hit patterns of single colored spots. The low molecular weight fragment library compounds, in contrast, were spotted in triplicates and therefore triplicate signals of αSyn interacting fragment hit compounds can be observed (Fig. 1b panel 1). Each array was assayed in duplicate during the screen. Fig. 1b panels 3 and 4 show scattergrams that illustrate the reproducibility of the raw SPR signals for each spotted sample in two independent repetitions for a representative of each type of microarray (the same representative microarrays shown in panels 1 and 2). The reproducibility of these subsets of compounds on the chosen microarrays is representative of results obtained for the entire library. Averaged SPR signals from replicate samples were filtered to remove outliers and JARRAY, NovAliX’s proprietary software routine, guided hit selection.

The screen identified 563 immobilized hit compounds interacting with αSyn, corresponding to a hit rate of 0.49%. The signal strength and hit rate generally were lower than that of previously screened globular proteins, but comparable to those obtained for our screen on the tau protein with HT-CM-SPR^23^. While for globular proteins of similar molecular weight to that of αSyn, signal to noise ratios of more than 10 were regularly obtained, for αSyn the signal to noise ratios ranged between 2 and 5. Nevertheless, as for the tau screen, hit compounds could be determined. Overall, hits included compounds which could be grouped into various compound classes and a number of structurally independent singleton compounds. We selected 152 representative hits out of the initial set of 563, based on parameters such as physicochemical properties, molecular weight, SPR signal strength, structural diversity, chemical tractability and overall attractiveness for drug development. This selected set contains structurally diverse and different lead-like and fragment-like compounds. Interestingly, a high number of positively charged and structurally different amines were identified. The molecular physicochemical properties indicate tractability for drug development of this compound set (Table 1 and Table S1). For further analysis of untethered hit compounds in follow-up assays, a subset of 65 compounds was selected for synthesis. This subset, the properties of which can also be found in Table 1, was chosen to cover chemical space of each structural class identified from the screen including singletons. All 65 compounds were synthesized devoid of the chemtag linker moiety, which was replaced with appropriate atoms/groups.

**Table 1.**
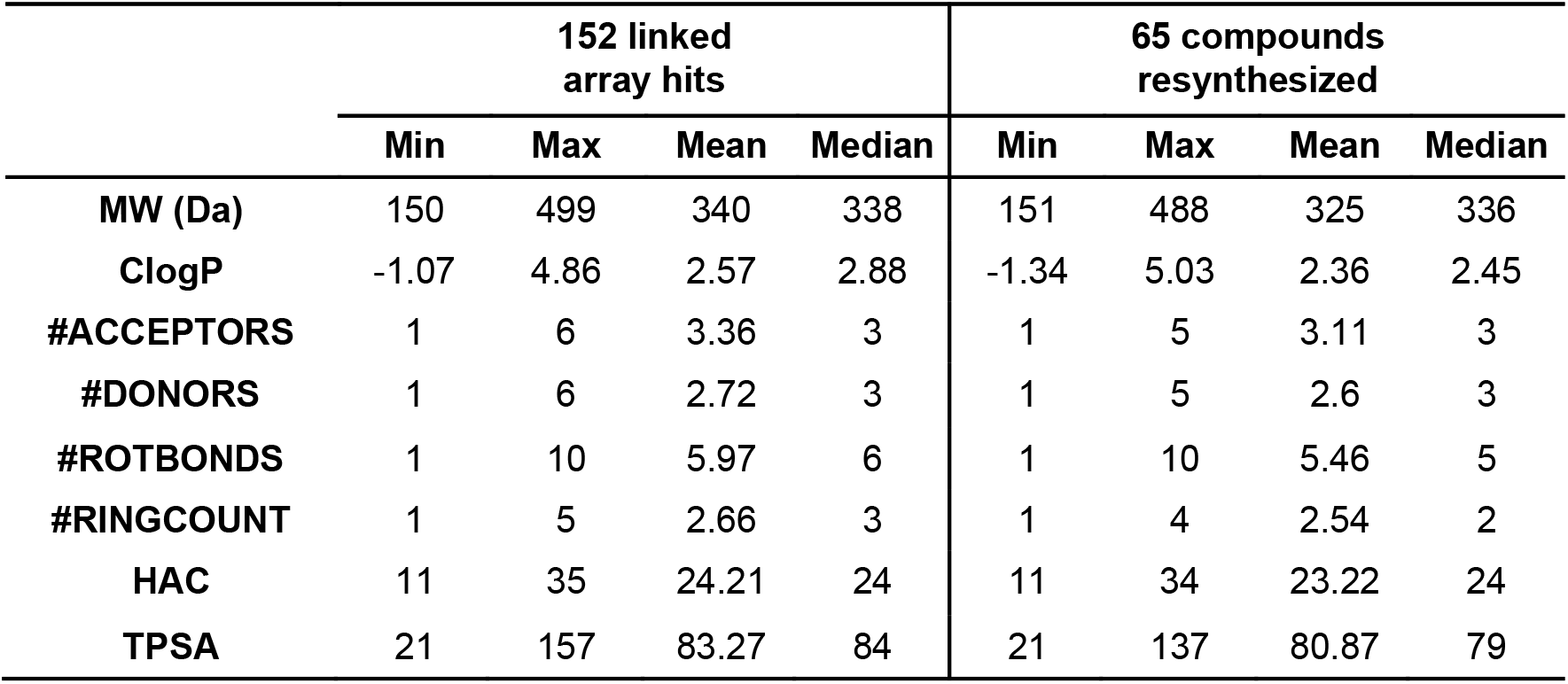
Physico-chemical properties of hit compounds. Analysis of the physico-chemical properties for the 152 representative hit compounds selected from the initial pool of 563 hits identified by the HT-CM-SPR screen of monomeric αSyn, as well as for the 65 resynthesized compounds. For immobilization to the chip surface all library compounds were tethered to the chip via a linker group R. In order to calculate the ClogP, H-bond acceptor count, H-bond donor count, the count of rotatable bonds and TPSA (topological polar surface area) for the 152 array hits, this R-group was virtually replaced by a carbon. For the calculation of the molecular weight (MW) and Heavy Atom Count (HAC) the R group was replaced by a hydrogen atom. For the 65 resynthesized compounds, the linker group R at the attachment point at the compound was synthetically replaced by other groups such as methyl groups, which are included in the calculation of all properties.

### Multiple hit compounds inhibit αSyn fibril and oligomer formation and one blocks cellular neurotoxicity

Misfolding of αSyn can result in the formation of toxic oligomers and amyloid fibrils^24^. Binding of small molecules to αSyn could block or enhance the transition of αSyn to β-sheet rich soluble oligomers and or fibrils depending on what conformation or sites are bound. We therefore screened the synthesized hit compounds in an αSyn aggregation assay, in which the fibril formation of the protein is monitored by Thioflavin T (ThT) fluorescence. Two compounds (576755 and 582032) are shown here which inhibit the fibrillization of αSyn (Fig. 2a). Shown in Fig. 2a for these compounds are examples of three separate experiments with the average of quadruplicate samples at each time point. Statistical analyses as described in Supplementary Information were used to determine significance of differences between compound and DMSO control. For the experiments summarized in Fig. 2a a significant difference from DMSO control was found for compound 576755 in 3 out of 3 experiments, and for compound 582032 in 2 out of 3 experiments with a trend to inhibition in the third experiment. Addition of compounds to αSyn monomer and ThT in the presence of preformed αSyn fibrils (Fig 2b, seeded fibrillization) demonstrates that compound 576755 blocks fibril growth in this rapid and robust paradigm showing activity as early as 2 hours in the incubation (Fig. 2b, right panel). The slower time-course of the traditional (non-seeded) fibrillization assay run in parallel is shown in Fig. 2a. Compound 576755 shows near half-maximal anti-fibrillization activity at a substoichiometric concentration of 25 μM (Fig. 2c) in the traditional fibrillization assay, in which αSyn is incubated at 70 μM.

**Figure 2.**
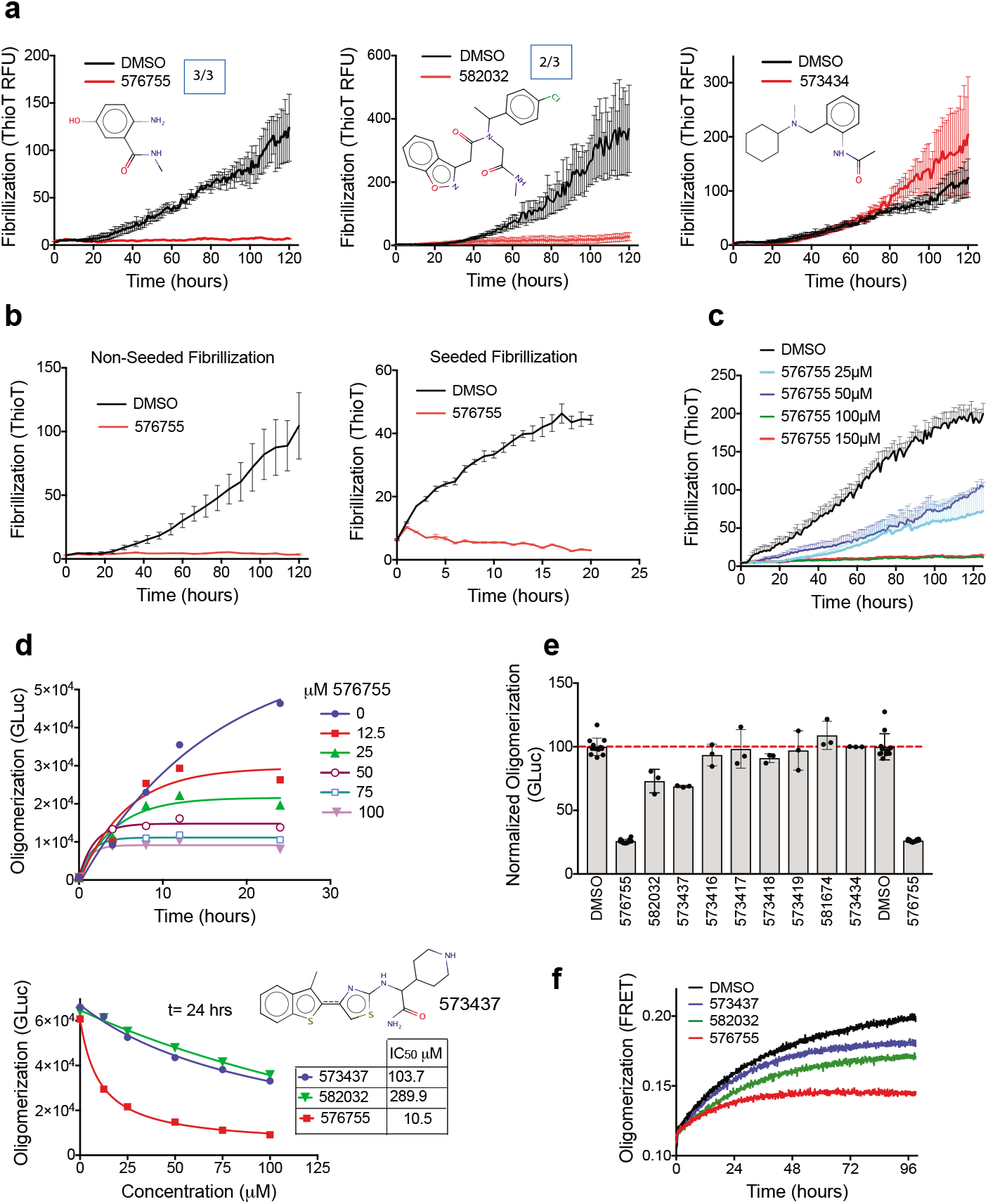
Multiple HT-CM-SPR screening hit compounds block αSyn misfolding. a) The impact of compounds on αSyn fibrillization activity, monitored by thioflavin T fluorescence, is shown. Examples are shown of three independent experiments for compounds tested at 300 μM in the αSyn fibrillization assay along with 0.5% DMSO control. Boxed numbers indicate the ratio of experiments demonstrating significant impact. Statistical analyses, (see Supplementary Information) indicate significant differences (p-value < 0.05) from DMSO for 576755 in 3 out of the 3 shown experiments, and for 582032 in 2 out of 3 experiments. Compound 573434 has no impact on αSyn fibrillization in a biochemical assay. **b)** Anti-fibrillization activity of compound 576755 at 300 μM is demonstrated in both the standard (left) and seeded-aggregation (right) assays in which αSyn protofibrils are added at the initiation of the assay. **c)** A dose response for compound 576755 in the fibrillization assay shows activity as low as 25 μM. For all fibrillization figures each data point represents the mean ± SEM of quadruplicate wells. **d)** Dose response of anti-oligomerization activity of three compounds are shown in a split Gaussia luciferase (GLuc) complementation assay executed as described in Methods. GLuc units are in arbitrary light units. The top panel shows the time course of inhibition by 576755 at multiple doses and the bottom panel shows the dose response and IC50s of all three active compounds at t=24 hours. **e)** The impact on oligomerization for all compounds discussed in this manuscript are shown. GLuc units are normalized to the mean of DMSO control. Means ± SD. are shown. **f)** The antioligomerization activities of the three compounds identified in panel **d** are validated in a FRET based αSyn oligomerization assay executed as described in Methods. Compound structures are also shown in Supplementary Fig. S4.

Compelling evidence supports pre-fibrillar αSyn oligomers as the pathogenic species in disease. αSyn oligomers are directly toxic to cells^25^, and mutations enhancing αSyn oligomer formation increase αSyn toxicity^26,27^. We therefore screened our compound set for impact on biochemical and cellular αSyn oligomerization. Two novel biochemical assays were developed for αSyn oligomerization based on either bioluminescent complementation of Gaussia luciferase (GLuc) split tags or on FRET of small molecule tags (see Supplementary Fig. S5 and Supplementary Information). The oligomerization assays we report here are sensitive, quantitative and reproducible assays of early αSyn misfolding and the bioluminescent assay in particular has a good dynamic range (Supplementary Fig. S5). The 65 synthesized hit compounds were tested in the biochemical bioluminescent complementation assay at 100 μM and three active compounds were identified (Fig. 2e), including the two compounds active in the fibrillization assay (Fig 2d: 576755, IC_50_ of 10.5 μM, and 582032, IC_50_ of 290 μM) and an additional compound (573437, IC_50_ of 104 μM). The activities of all three compounds were verified in the FRET based αSyn biochemical oligomerization assay (Fig 2f), demonstrating that the compound action was directed at αSyn and not the luciferase tags. The development of the novel oligomerization assays, which are much less variable than fibrillization assays, allowed us to identify the additional compound 573437 which most likely due to the variability of the fibrillization assay did not reach significance during the fibrillization testing.

Maximal concentrations of compounds not showing toxicity were tested in H4 neuroglioma cells for impact on αSyn cellular oligomerization using a bioluminescent protein-fragment complementation assay^28^. The only compound that reproducibly reduced cellular oligomers was 576755 (Fig. 3a). Western controls show no significant change of cellular αSyn protein levels by 576755 (Fig. 3b, a trend to increase is seen). To test the effects of 576755 on αSyn-induced neurotoxicity, we used a primary midbrain culture model of PD^29^. As a control, we first established that there is no detectable toxic effect on primary dopaminergic neurons exposed to 60 μM 576755 alone (Fig. 3c). Transduction of the primary cultures with an adenovirus encoding A53T αSyn, a PD-linked genetic mutant found to form toxic oligomeric species more readily than the wild type protein^5,30^, results in a 30–40% reduction in the number of TH-immunoreactive neurons (Fig. 3d). In contrast, transduction with a virus encoding the control protein LacZ has no effect on neuron viability^31^. Treatment of the primary midbrain culture with 576755 at 60 μM, suppressed A53T αSyn neurotoxicity, and a trend towards a similar inhibitory effect was observed for cultures treated with the compound at 10 μM (Fig 3d).

**Figure 3.**
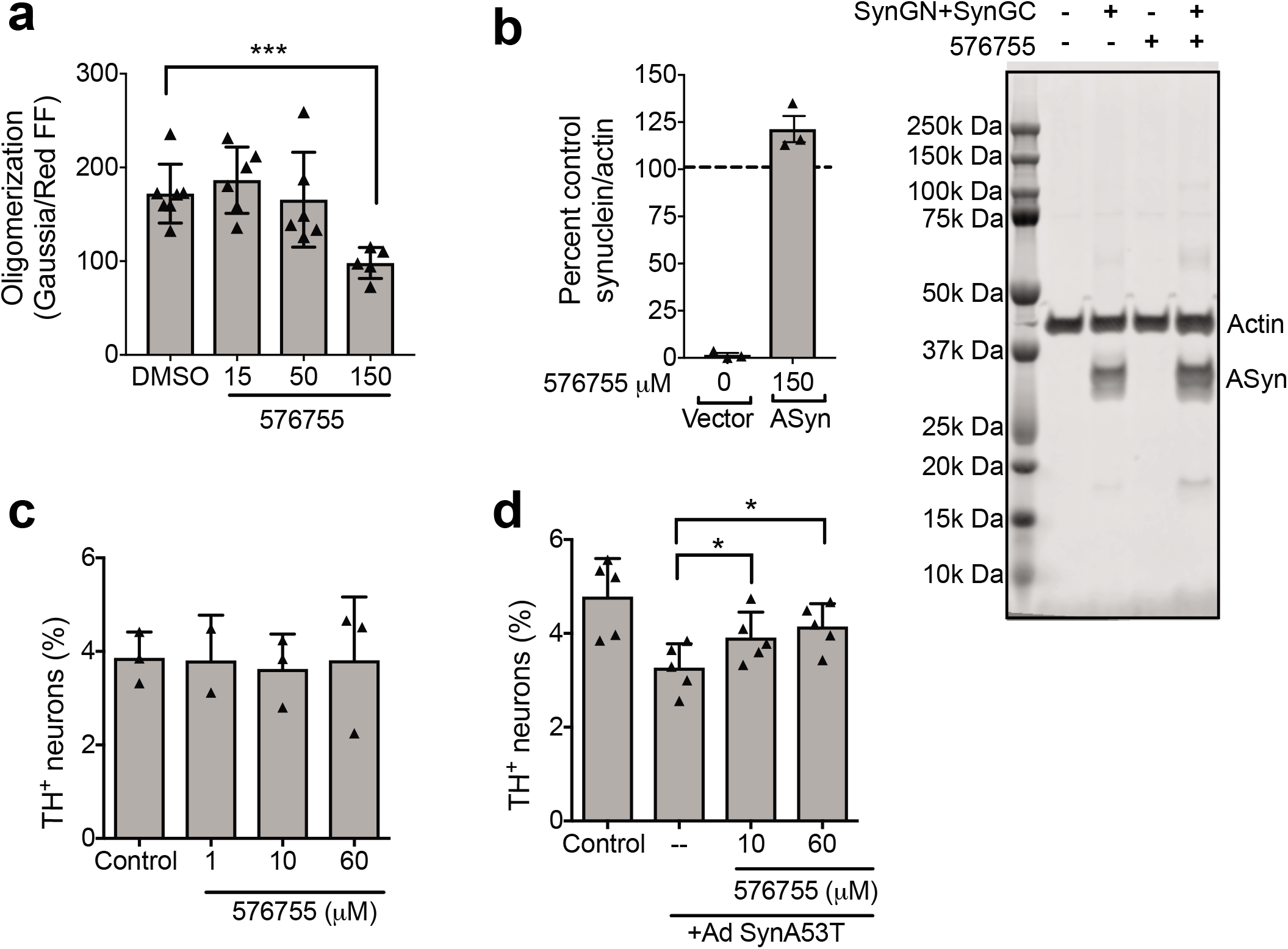
Cellular activities of anti-aggregation compound 576755. **a)** 576755 reduces αSyn oligomerization in H4 cells as measured by complementation of αSyn proteins with split luciferase tags executed as described in methods. Trace firefly luciferase co-transfected provides a normalization measure for transfection efficiency. 0.3% DMSO is present in all samples. Data are plotted as means +/- SD. Representative results from 3 independent experiments. A reduction in cellular oligomerization by 576755 was determined by one-way ANOVA with Dunnett’s post-test. **b)** 576755 does not reduce on αSyn levels in H4 cells as determined by Western analyses (trend towards small increase seen). Right) representative Western from one experiment of a merged image detecting actin and αSyn from H4 cells transiently transfected with vector control or αSyn containing N or C terminal split luciferase tags (SynGN+SynGC) and treated with 0.3 percent DMSO or 150 μM 576755. The two αSyn bands correspond to different tags. Entire length of blot shown. Outline of full blot shown by black lines. Left) αSyn was quantitated in 3 separate biological replicates. Samples were normalized to cells transfected with αSyn without drug treatment to allow comparisons between blots. There is no significant difference in αSyn levels in 576755 treated vs. untreated cells (100) as determined by one sample t test. Data are plotted as mean ± SD. **c)** 576755 is not toxic and **d)** alleviates loss of dopaminergic neurons induced by the A53T mutant of αSyn. Primary rat embryonic midbrain cultures were nontransduced or transduced with adenovirus encoding A53T αSyn (+Ad SynA53T), in the absence or presence of 576755. The cells were then stained immunocytochemically for MAP2 and TH. Preferential dopaminergic cell death was assessed by evaluating the percentage of MAP2-positive cells that also stained positive for TH. Data are plotted as the mean ± SEM. n=2-3 for the neuron toxicity analysis and n=5 for the reversal of αSyn toxicity. Shown are representative results from 5 independent experiments. Statistics used a oneway ANOVA with Tukey post-test after square root transformation of the data. *p < 0.05 where shown. ***p < 0.001. **** p < 0.0001. ns is not significant.

### Multiple hit compounds reverse αSyn mediated inhibition of vesicular dynamics

Elevated αSyn can interfere with the creation, localization, and/or maintenance of vesicle pools^32,33^. We have demonstrated that αSyn overexpression impairs phagocytosis in H4 neuroglioma cells, in primary microglia isolated from αSyn transgenic animals overexpressing αSyn and in cells from human PD patients by blocking recruitment of vesicular components necessary to support a phagocytic response ^34^. We further reported that an αSyn binding compound derived from an *in silico* screen, 484228, restored normal phagocytosis in the face of ongoing αSyn overexpression^20^. We thus tested whether the αSyn binding hit compounds found here could protect against this αSyn over-expression induced dysfunction. The synthesized hit compounds were screened first for cellular toxicity at ranges from 1 μM to 100 μM. 53 compounds, showing no toxicity at 1 μM, in the absence of serum were subjected to further screening for their impact on phagocytosis in H4-neuroglioma cells overexpressing αSyn^20,34^ at two concentrations below that for which toxicity was seen. Seven compounds showed robust activity in reversing αSyn mediated inhibition of phagocytosis in H4-neurogloma cells in the initial screen. Five of these compounds were retested at 10 and 1 μM and 100 nM concentrations (Fig. 4a). All compounds showed complete reversal at 1 μM and two compounds (573418, 573419) showed complete reversal at 100 nM (Fig. 4a). Western analyses demonstrate that none of these compounds alters αSyn protein levels (Fig. 4b) and none of these compounds impaired αSyn oligomerization (Fig. 2e).

**Figure 4.**
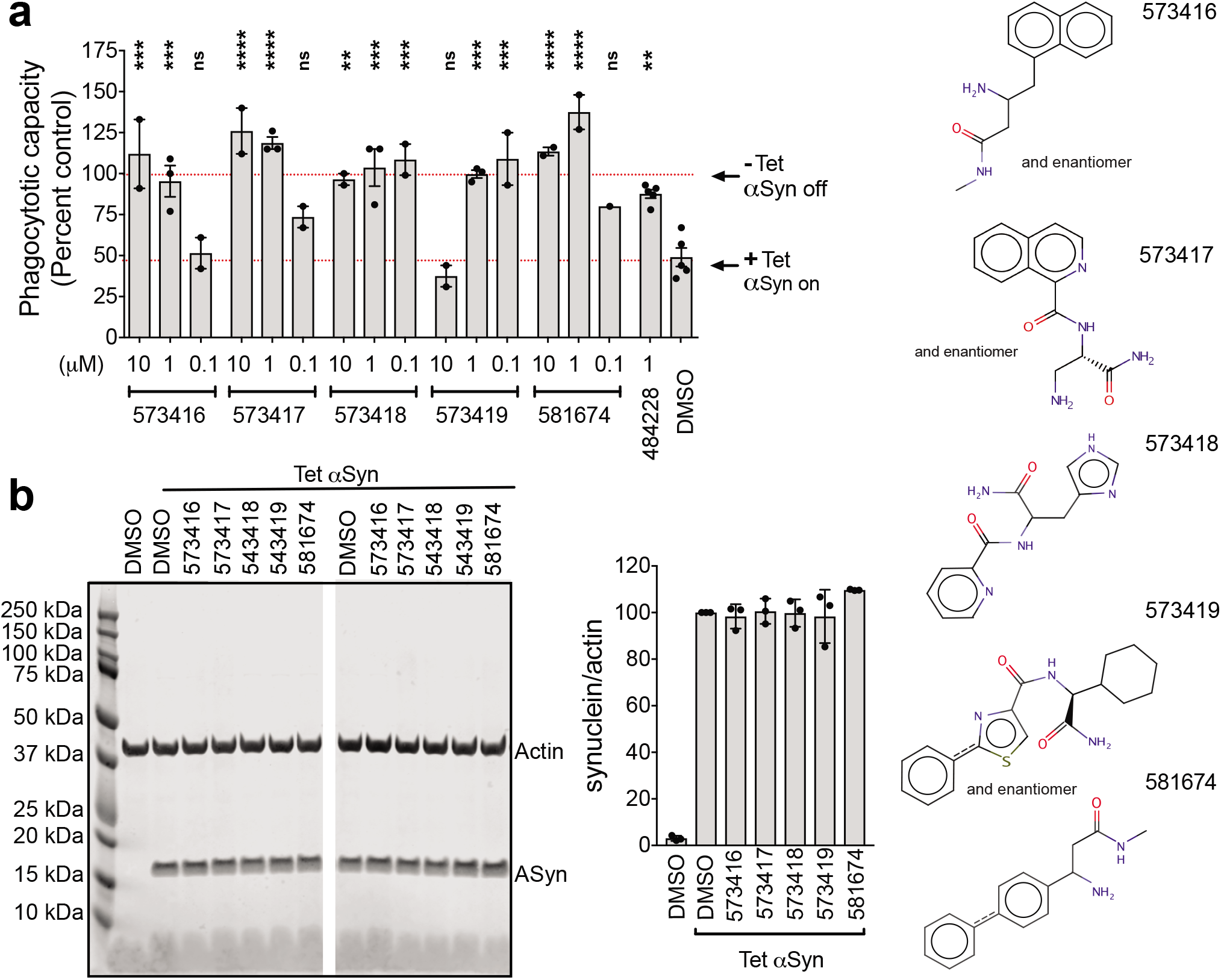
Multiple HT-CM-SPR screening hit compounds alleviate αSyn mediated inhibition of phagocytosis in H4-Neuroglioma cells: H4 neuroglioma cells over-expressing αSyn from a tetracycline-inducible promoter were cultured for 24 hours with compound in the absence or presence of αSyn induced by tetracycline. After 24 hours of induction cells were a) fed 4 micron beads for 90 minutes and a phagocytic index was measured by quantitating the amounts of engulfed beads on an imaging reader or b) analyzed by Western blot to determine αSyn levels. a) The phagocytic capacity was calculated by normalizing the indicated samples to the phagocytic capacity of un-induced cells not overexpressing αSyn (Tet off). Each point corresponds to a separate experiment denoting the average of 6 well replicates. Means ± SD for the combined multiple experimental averages are shown. Control compound is 484228, identified in a prior *in silico* screen^20^. n=2 or 3 different experiments as shown. Significance was determined by ANOVA with Dunnet’s correction. *p < 0.05, *** p < 0.001, **** p < 0.0001. ns is not significant. b) Cells were untreated or treated with tetracycline to induce αSyn and treated with DMSO or compounds and analyzed by Western blot for actin and αSyn levels. Left: Representative Western blots of merged actin and αSyn signals of individual wells treated with DMSO or compounds. Image cut as shown to remove irrelevant samples. Entire length of blot shown. Outline of full blot shown by black lines. Right: Westerns from multiple replicates were quantitated and the αSyn band intensity was normalized to that of actin. Each data point is a separate well. No compounds showed significant impact on αSyn levels. Compound structures are also shown in Supplementary Fig. S4.

### One hit compound blocks αSyn cellular transmission without apparent impact on misfolding

Lower-order αSyn oligomeric species and fibrils propagate their misfolded structure and transmit from cell-to-cell in a prion-like manner^35–37^. We therefore tested selected compounds in a cellular model of αSyn transmission^38,39^. Figure 5 shows that compound 573434 retards αSyn transmission both in B103 rat neuroblastoma cells using a co-culture paradigm (at the Gladstone Institutes; Fig. 5a) and as executed at a different institution in primary neurons using a separate chamber culture paradigm (at UCSD; Fig. 5b and 5d) with activity at 10 μM 573434. 573434 has no observed impact on αSyn fibrillization nor oligomerization (Fig. 2a and 2e) and does not impact αSyn levels in the donor cells as measured by quantitative immunohistochemistry (Fig. 5c) and by Western analyses (Supplementary Fig. S6). Furthermore 573434 does not impact the secretion of αSyn from cells as determined by Western analyses (Fig. 5e). Thus, the mechanism of action of 573434 is not due to direct action on αSyn misfolding, nor on expression or excretion of αSyn.

**Figure 5.**
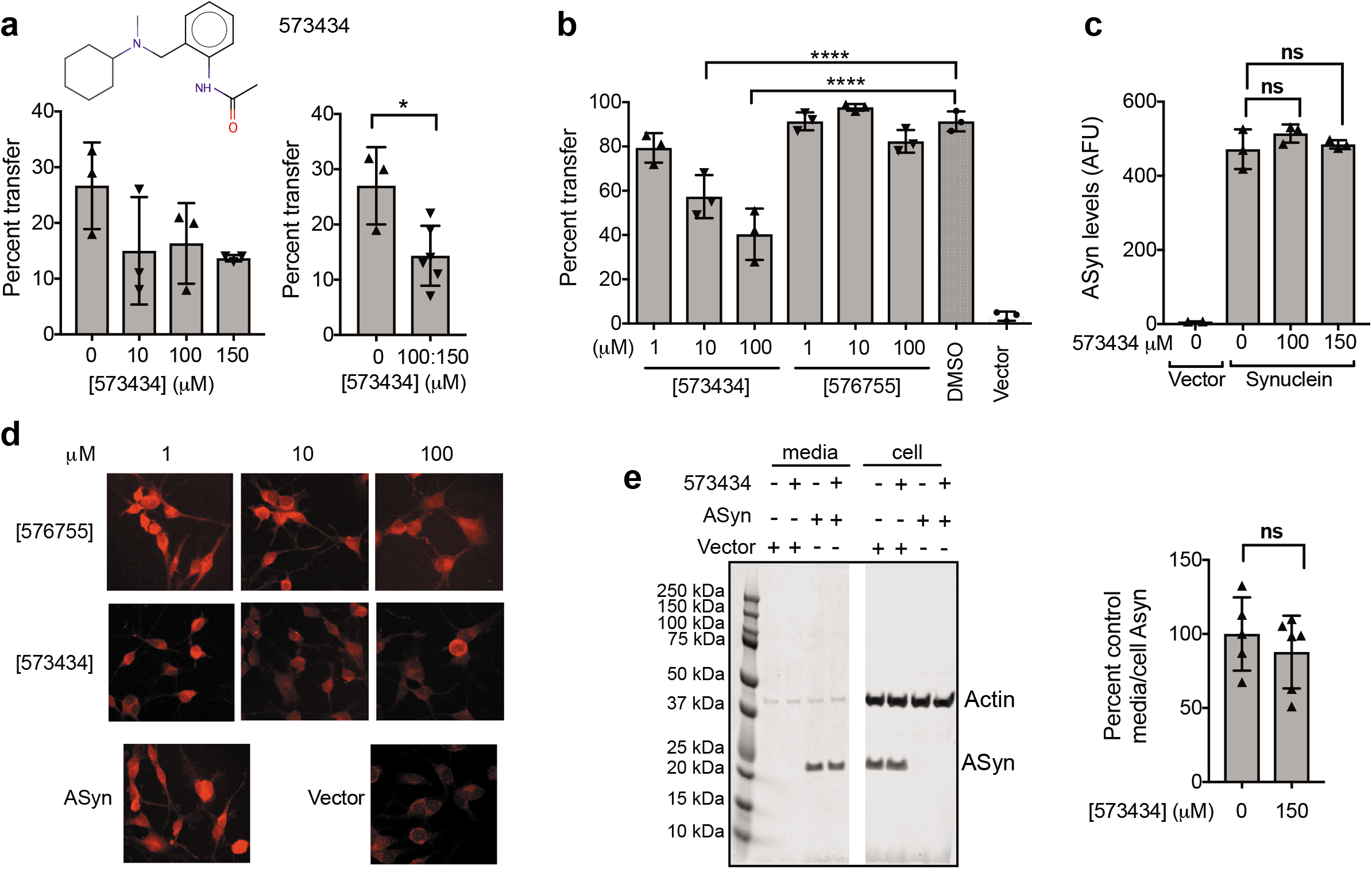
One HT-CM-SPR screening hit compound blocks cell-to-cell transmission of αSyn. **a)** The impact of 573434 on transmission was tested on co-cultured donor and recipient B103 neuroblastoma cells and mean percentage of receiving cells with αSyn is shown. The same data are shown separating all compound doses (left) or combining 100 and 150 μM (right). 3 coverslips/well per group, 10-15 pictures per coverslip, and 43-168 cell total per group were analyzed. Combined 100 and 150 μM drug samples give statistically significant differences from untreated by unpaired t test (n=3). **b)** Donor and acceptor primary neurons were cultured in separate chambers of a Transwell and αSyn in acceptor cells was measured after treatment with 573434 or 576755. Vector control is lentiviral vector not expressing αSyn. 10 and 100 μM 573434 retarded transmission whereas 576755 had no impact as determined by ANOVA with Dunnett’s correction. **c)** Total αSyn levels were determined from the fluorescent intensity from αSyn antibody (arbitrary fluorescence units, AFU) in the images used in panel a). 3 coverslips per group, 8-11 pictures per coverslip, and 24-54 cell total per group were analyzed. 573434 does not impact overall expression of αSyn as determined by unpaired t test. **d)** Representative images of αSyn transmission in primary neurons treated with 573434 and 576755. **e)** Cells and media produced by B103 cells infected with control or αSyn-lentivirus and treated with DMSO or 150 μM 573434 were analyzed by Western blot for actin and αSyn levels. Left: Western blot of both actin and αSyn proteins of duplicate wells. Image cut as shown to remove irrelevant samples. Entire length of blot shown. Outline of full blot shown by black lines. Right: the ratio of αSyn in the media to αSyn in the cell was quantitated from three separate experiments with duplicate wells (n=6). αSyn levels in cells and media were normalized to actin levels in cells of that well and then further normalized to control (DMSO) sample on the same gel to allow for comparison between experiments. 573434 has no impact on the amount of αSyn secreted into the media. For all figures drug treated and control cells are in equivalent levels of DMSO (0.15 to 0.3%). Each symbol represents a singe coverslip in a unique well. Mean ± SD. *p < 0.05, ***p < 0.001, ****p < 0.0001, ns–not significant. Compound structure is also shown in Supplementary Fig. S4.

## Discussion

In a prior study we reported on a novel strategy to target the monomeric intrinsically disordered ensemble of αSyn by using a structure-based computational docking screening approach to identify small molecules predicted to bind to monomeric αSyn, and then testing their impact in diverse αSyn mediated biochemical and cellular assays. An advantage of this approach is its potential to identify compounds that have a variety of effects related to αSyn malfunction or misfolding. This effort identified one compound, 484228, which reversed the impairment of phagocytosis, dopaminergic neuronal loss and neurite retraction caused by overexpression of αSyn^20^.

In order to expand upon this strategy, we applied a biophysical-based binding screen that detected the interaction between a small molecule and native monomeric αSyn, using SPR technology, an approach we have successfully used to identify anti-aggregation compounds for the tau protein^23^. The current screen identified over 500 hundred small molecules interacting with αSyn. We further showed that some of these compounds could beneficially impact αSyn malfunctions observed not only in aggregation assays, as we had found for the previous tau protein screen^23^, but also in multiple cellular malfunction assays. Our anti aggregation compound 576755 blocked cellular αSyn oligomer formation and protected dopaminergic neurons against αSyn toxicity. We found compounds similar to our reported 484228^20^ that reversed the impairment of phagocytosis by αSyn, and we also found a different compound that blocks cell-to-cell transmission of αSyn. Further studies will be needed to establish whether these compounds interact with the different conformations of monomeric αSyn and/or higher-order assemblies. Overall, however, as we had anticipated, an expanded screen for compounds binding to monomeric αSyn yielded a variety of novel classes of compounds having different effects on αSyn mediated pathological processes providing a rich starting point for drug-discovery.

There is ample evidence that links αSyn misfolding and aggregation to the onset and progression of PD^40^. It is still unclear, however, which conformations of misfolded αSyn contribute most to pathogenicity. Prior screens for compounds modulating αSyn aggregation have relied largely on screening in fibrillization assays, and many have yielded compounds that block aggregation in a nonspecific manner mediated through polymeric self-stacked forms of the compounds^41,42^ or via reactive quinone formation^43^ which complicates translation into effective drugs. Although our most potent anti-aggregation compound, 576755, has quinone potential, two additional misfolding blocking compounds without this liability were found in this screen using novel oligomerization assays. The biochemical bioluminescent protein complementation αSyn oligomerization assay, in particular, is a highly useful and novel assay in that it is amenable to high-throughput screening and does not rely on shaking or detergents to accelerate αSyn misfolding as is the case for a recently published αSyn high-throughput oligomerization assay^44^. The ability to identify anti-aggregation compounds through a fundamentally different screening paradigm - that of SPR-based screening using monomeric αSyn - allowed us to obtain alternative compounds inhibiting aggregation. This will potentially expand our repertoire of anti-aggregation compounds to include compounds of possibly novel and druggable mechanisms.

The role of αSyn in modulating vesicle dynamics in cells is well established^33,45–48^, and there are links between αSyn toxicity and vesicular dysregulation^33^. We have reported that phagocytosis, which involves both mobilization and extrusion of vesicles, is impaired by αSyn over-expression in cultured cells and *in vivo* in transgenic mice as well as in cells from Parkinson’s patients^34^ suggesting that this dysfunction is a relevant target for therapeutic intervention in PD. Our experience using 484228 as a control compound in scores of assays at high doses is that it never restored phagocytosis inhibited by αSyn overexpression beyond 80 percent of that in cells with endogenous αSyn levels. The compounds reported herein restore phagocytic capacity to 100 percent of control and thus are likely to act via mechanisms distinct from that of 484228. Thus, the identification of multiple drug-like compounds with different scaffolds reversing αSyn impairment of phagocytosis expands upon our earlier results with compound 484228.

A central role of transmission and propagation of misfolded αSyn in disease is widely accepted^37^. Although antibodies to αSyn are being tested in the clinic to block pathogenic spreading^49^, small molecules targeting IDP spread are limited to aggregation inhibitors and to a single pharmacological chaperone^50^. The anti-fibrillization compound 576755 did not impact transmission, despite its ability to reduce cellular oligomers. This could be due to differing drug sensitivities in the cells used in these assays. Nevertheless, we have identified a different small molecule, 573434, which blocks the transmission of αSyn from one neuronal cell to another. This compound does not change overall αSyn levels, ruling out a nonspecific impact on the vector. It furthermore does not impact the overall levels of αSyn secreted from the cells. In biochemical assays it impacts neither αSyn oligomerization nor fibrillization indicating a novel mode of action for this molecule. There remain a number of possible mechanisms including 1) reduction of overall uptake of αSyn into receiving cells or 2) reduction of levels of a subpopulation of αSyn that is preferentially taken up in receiving cells. For example, Danzer and colleagues have shown that only the fraction of extracellular αSyn that is packaged into exosomes in the media is transmitted into receiving cells^51,52^. Levels of this specific αSyn pool could be reduced by 573434. Further studies are needed to elucidate an exact mechanism of action for this compound.

The high-throughput approach described here may also be applied to other IDPs. Indeed, in a related study, we applied HT-CM-SPR screening for small molecules interacting with monomeric full-length tau, which resulted in the discovery of a variety of compounds which were able to reduce the aggregation of tau *in vitro* and in a cell model^23^. Hence, these studies add to the accumulating evidence that small molecules can bind to IDPs, such as αSyn^20^, tau^23,53^ and the Aβ peptide^54^, despite their overall lack of persistent 3D structures.

In conclusion, these results support the notion that the dynamic monomeric solution state of αSyn can be successfully targeted by drug-like small molecules to reverse PD relevant dysfunctions. Our identification of small molecules impacting multiple types of αSyn malfunction demonstrates that targeting αSyn with screens employing ensembles of predominantly monomeric protein offers a rich and effective opportunity for drug discovery. We thus anticipate that this approach will be useful for the development of small molecule therapeutic candidates for Parkinson’s disease and other diseases associated with IDP misfolding.

## Methods

Additional details are in Supplementary information.

### SPR Screening of Monomeric αSyn

The binding of αSyn was evaluated on a set of 96 control ligands of defined physical chemical properties in order to optimize buffer composition, ligand surface density and αSyn concentration. Minimum background and maximal total signal were obtained using 900nM αSyn protein in 25mM Tris and 100mM NaCl, pH 7.4. αSyn at 10x that of screen concentration was predominantly monomeric as measured by DLS (>99.9%, see Supplementary Information). Freshly diluted αSyn in screening buffer was centrifuged through a 100kD cut-off filter, before incubation at room temperature on the arrays for up to 3 hours. Duplicate chips were run, with unique samples for the lead-like library and triplicate samples for the fragment library. Multiplicate SPR signals were averaged per compound. Compounds with standard deviation more than 0.5, or for which the purity before spotting or the saturation per spot was less than 80 percent were discarded.

### Fibrillization assays

Prior to each aggregation assay, purified αSyn at 5 mg/mL was treated with 6M guanidine and dialyzed 36 hours in a 3500 MW cutoff Slide-A-lyzer with three changes of 20 mM sodium phosphate and 100 mM NaCl pH 7.4 buffer. αSyn was centrifuged at 130,000g for 40 minutes in a Beckman Ultracentrifuge to remove seeding species and supernatant used for assays. For initial testing, compound was diluted to appropriate concentrations in 0.5% DMSO and controls contained equivalent DMSO concentrations. Assay samples contained 20 μM Thioflavin T, filtered through a 0.45 μm Acrodisc filter prior to each assay, 70 μM αSyn in 20 mM sodium phosphate and 100 mM NaCl pH 7.4. One teflon bead, (0.125” PTFE, Grade 2 polished) was in each well of a 96 well black with clear bottom assay plate (Corning) in a total volume of 500 μL. The plate was sealed with Mylar and parafilm, placed at 37°C in a Tecan F200Pro Plate Reader and shaken at an amplitude of 6 mm for 120 hours continuously except for a brief time during reading of the plate. Thioflavin T fluorescence was detected by excitation at 440 nm and reading emissions at 485 nm each 60 minutes throughout the assay. To generate αSyn seeds for the seeding assays, αSyn at 10 mg/mL was incubated with stirring at 43°C for 24 hours followed by exhaustive sonication. 1% (by mass) of seeds were added to the fibrillization assay. Samples were plated in quadruplicate, and the replicates averaged for each experiment for plotted data. Data were analyzed in Excel using XL-fit sigmoidal model #600.

### Biochemical αSyn oligomerization assays

For the split Gaussia luciferase (GLuc) oligomerization assay 10 μM each of αSyn-GLuc1 and αSyn-GLuc2 (see Supporting Information: N-terminal or C-terminal fragment of GLuc fused to the C-terminus of αSyn) were incubated in a buffer containing 1 mM MgCl_2_, 1 mM ADP, 50 mM Na_3_PO_4_, 0.02% NaN_3_ and 1 mM DTT at pH 7.4 at 37°C in a thermal cycler (BioRad DNA Engine PTC-200). In some cases, when specified, the samples were incubated with shaking in an incubator at 37°C. At 0, 6, 12, and 24 hours of incubation GLuc activity was quantified using the BioLux Gaussia Luciferase Assay Kit (NEB) according to the manufacturer’s instructions with luminescent detection on a SpectraMax M5 Microplate Reader (Molecular Devices) using a 5 minute delay after the addition of substrate and 3 seconds of integration time.

For the FRET oligomerization assay,10 μM each of αSyn-Q-(Position#)C-Cy3 (donor) and αSyn-Q-(Position#)C-Cy5 (acceptor) in PBS (pH 7.4) were dispensed into black/clear bottom 384-well plates (Greiner) using a Mantis liquid handler (Formulatrix). The plate was sealed with AbsorbMax film (Sigma-Aldrich) and incubated at 37°C. The FRET between donor and acceptor was measured on a SpectraMax M5 Microplate Reader (Molecular Devices). An excitation wavelength of 525 nm for αSyn-Cy3 (donor), and emission wavelengths of 570 nm and 670 nm for αSyn-Cy3 (donor) and αSyn-Cy5 (acceptor) respectively were used. The plate reader PTM sensitivity was set to medium and assays executed without mixing. The signals were read at 10 minute intervals for 99 hours at 37°C. Efficiency of FRET was calculated as follows: EFRET=I_Acceptor_/I_Donor_, in which I_Acceptor_ is acceptor emission intensity, I_Donor_ is donor emission intensity.

### Cellular αSyn oligomerization assay

H4 neuroglioma cells (HTB-148; ATCC) were passaged in DME containing 10% fetal calf serum (FCS). They were plated into 10 cm dishes at 7.5×10^5^ cells per dish and the following day transfected with DNA and Fugene (Promega, Madison, WI) according to the manufacturer’s directions at a ratio of 3:1 with 7 μg each of Syn-Luc1 (S1) and Syn-Luc2 (S2) plasmids^55^ (also referred to as syn-hGLuc(1) and syn-hGLuc(2)^55^ respectively, kind gifts from Dr. Pamela McLean) and 3 μg of red firefly expressing plasmid (pCMV-Red firefly Luc, ThermoFisher, Waltham, MA) per dish. S1 and S2 plasmids express αSyn containing split Gaussia luciferase with either the N-terminal fragment (S1) or C-terminal fragment (S2) attached to the C-terminus of αSyn. After 24 hours, the DNA containing media was removed and cells were fed fresh DME media containing 10% FCS. The following day, cells were plated into polyD-Lysine coated clear bottom white well 96 well plates at 1.5×10^4^ cells per well in Opti-MEM without phenol red with penicillin and streptomycin. After 4 hours cells are treated with drugs and incubated for 20 hours. Gaussia luciferase activity was measured in the dark 0.1 sec after injection of 100 μl/well of 40 mM coelenterazine, substrate (NanoLight, Pinetop, AZ) using a 2 second integration on a Veritas Microplate Luminometer (Promega, Madison, WI). The cells were subsequently assayed for red Firefly activity using the ONE-GloTM Luciferase Assay System kit (Promega, Madison, WI) on a Spectramax M5 (Molecular Devices, San Jose, CA) plate reader as a normalization measure. 100 μl of media were also assayed for red firefly activity. The impact of compounds on H4 cell toxicity was assayed using the CytoTox-Glo kit (Promega, Madison, WI).

### Phagocytosis assay

A human neuroglioma H4 cell-derived cell line stably over-expressing αSyn from a tetracycline inducible promoter^20^ was grown in serum-free X-VIVO media (Lonza Group, Basel, Switzerland). Cells were seeded in 96-well plates at 100K cells per well. The next day, compound was added along with 5μg/ml tetracycline to induce αSyn overexpression. Cells were cultured overnight and the next day were fed 4 μM red fluorescent beads (In Vitrogen, Carlsbad, CA) for 90 minutes at a cell-to-bead ratio of 1:10. Plates were gently washed with 100μl/well media twice, fixed and stained with HEMA3 (Thermo Fisher, Waltham, MA). Plates were dried overnight and read on an ArrayScan (Thermo Fisher, Waltham, MA). As the HEMA3 stain absorbs light, the internalized beads are less fluorescent than the outside beads. Tet/non-tet and 484228 samples were run on each plate.

### Transmission assay

Lentivirus was made from the pLV-aSyn vector (LV-aSyn)^56^ expressing his tagged αSyn or control green fluorescent protein vector (LV-control) as described^38^. Titers were determined by P24 protein ELISA (primary neuronal experiments) or quantitative PCR of genomic DNA using transducing units of virus expressing green fluorescent protein titrated on HEK293 cells serving as a normalizing control (rat B103 experiments). Rat B103 neuroblastoma cells were grown in DMEM with 10% FBS at 37°C in 10% CO_2_.. Cryopreserved mouse cortical neural stem cells (MCNS) (MilliporeSigma, Burlington, MA) were grown according to manufacturer recommendations. αSyn transmission between cells was done using the methods described^38,39^ in one of two ways: 1) B103 neuronal cells: donor B103 neuronal cells plated in 6 well plates at 250,000 cells per well were incubated with LV-aSyn or LV-control at an MOI of 20 for 48 hours. B103 acceptor cells were labeled with the 595-Qtracker labeling kit (Life Technologies, Carlsbad, CA) according to the manufacturer’s protocol. Donor and acceptor cells were trypsinized and replated together in a 12 well plate at 50,000 cells/well each on polyL coated coverslips. 4 hours post plating drugs were added. After 48 hours of co-incubation cells were fixed in 4% paraformaldehyde (PFA). Coverslips were stained with Syn1 antibody (BD Franklin Lakes, NJ) at a dilution of 1/500 and imaged on a Zeiss LSM 880 laser scanning confocal microscope and the percentage of receiving cells positive (cutoff of 10 standard deviations above negative control) for αSyn calculated by using ImageJ. Total αSyn levels were quantitated on the same images quantitating total fluorescence intensity from the Syn1 antibody using ImageJ. 2) MCNS neurons: Donor MCNS neurons were infected with LV-aSyn at a MOI of 20 for 48 hours, trypsinized and replated in 12 well cell culture inserts containing a 0.4 μm PET membrane (Thermo Fisher, Waltham, MA) at 50,000 cells/well. Acceptor MCNS were plated in the bottom chamber of the 12 well Transwell plate at 50,000 cells/well on a poly-l-lysine coated glass coverslip. The 0.4μm filter, allows passage of secreted proteins but restricts direct cellular contact. 4 hours post plating, drugs were added. Cells were co-incubated for 48 hours and fixed with PFA, followed by immunocytochemical analysis.

### Statistical Analysis Methods

Methods for the analyses of compound activity in the synuclein fibrillization assay are in Supplementary Information. Other statistical analyses were run using GraphPad Prism software as described. Multiple samples were compared using one-way ANOVA with Dunnett’s and an alpha of 0.05. In analyzing percentage cell viability data by ANOVA, square root transformations were carried out to conform to ANOVA assumptions, and Tukey’s post *hoc test* was used for multiple comparisons^31^. In cases where two samples were used t tests with an alpha of 0.05 were used. General practice significance nomenclature was used (0.1234 (ns), 0.0332 (*), 0.0021 (**), 0.0002(***), <0.0001(****) unless otherwise indicated.

### Compound analyses and handling

The 65 resynthesized compounds chosen for testing in functional assays were verified as the indicated structure and of sufficient purity by ^1^H-NMR and by liquid chromatography and mass spectrometry analyses (LC-MS). These 65 compounds were at least 85% pure, with most over 95% pure. The active compounds were all over 90% pure. Additional information including the LC-MS determined purity and ^1^H-NMR peaks for the 9 active compounds described herein are listed in Supplementary Information.

For all compound testing, controls are run in the equivalent amount of DMSO.

## Associated content

Supplementary Information includes detailed data with expanded description and expanded methods.

## Acknowledgements

This manuscript is dedicated to Dale Schenk, who was an inspiring and visionary leader^57^. We are grateful to Andrei Konradi and Al Garofalo for assistance in selecting compounds, to Paul Beroza and Ali Ozkabak for compound database management, to Kari Callaway for compound solubility testing and to Kari Callaway, to Karin Regnstrom and Donald Walker for helpful discussions. We are grateful to Pamela McLean and Simon Moussaud for the S1 and S2 plasmids and advice on developing the cellular oligomerization assay, to Anthony L. Fink and Katerina Levitan for assistance in developing the high throughput synuclein fibrillization methods to Alexander Buell for sharing his seeded synuclein fibrillization methodologies. The research carried out at NovAliX (originally Graffinity Pharmaceuticals) and the University of Cambridge was supported by Elan Pharmaceuticals. Research at UCSF and the Gladstone Institutes was supported by and the National Institute of Neurological Disorders and Stroke of the National Institutes of Health under award number R21NS092897 (LM). Additional support was provided by R01 grant NS049221 and by a grant from the Branfman Family Foundation (J.-C.R.). We are particularly grateful to the late T. Gary Rogers and the Rogers Family Foundation for their support of this work (LM, DAA). DAA is also supported by the Howard Hughes Medical Institute.

## Author Contributions

Designed and performed research, analyzed data on HTS screen (T.N., H-D.J., D.S., R.S., I.O., X-H.C., B.S., J.P.A, G.T.), designed and performed research, analyzed data on biochemical fibrillization testing (N.C., C.W.B. J.C., M.J., S.B. M.B., W.H.), designed and performed research, analyzed data on biochemical oligomerization testing (J.T., D.A.A. L.M.), designed and performed research, analyzed data on cellular oligomerization testing (A.B., P.N., M.T., J.T., L.M.), designed and performed research, analyzed data on neurotoxicity testing (M.T., J-C.R., B.S., E.M., L.M.), designed and performed research, analyzed data on phagocytosis testing (S.G., J.M., T.Y.), designed and performed research, analyzed data on transmission testing (A.B., B.S., E.M., L.M.), wrote the paper (L.M., G.T., D.A.A., J.C.) designed research, supervised and analyzed data on all aspects (L.M., G.T., D.S.).

All authors reviewed the manuscript.

