## Supplemental Information for "Novel Small Molecules Targeting the Intrinsically Disordered Structural Ensemble of α-Synuclein Protect Against Diverse α-Synuclein Mediated Dysfunctions"

***Quality control analyses of αSyn used for high-throughput chemical microarray SPR imaging (HT-CM-SPR) screening***

Gel electrophoresis, $^{1}H$-$^{15}N$ HSQC NMR and DLS analyses were performed on the αSyn sample used for screening to ensure the integrity, purity and monomeric nature of the protein used in the screen. Multiple aliquots of the purified protein were frozen for single use and used for all analyses and all screening. The protein ran as one peak during the size exclusion chromatography used in the final purification step (data not shown). Electrophoresis indicates that αSyn was pure and runs predominantly as a full-length monomer in the presence of sodium dodecyl sulfate (SDS) (Fig. S1). $^{1}H$-$^{15}N$ HSQC NMR analyses were used to compare the dynamic ensemble of the αSyn sample used for screening to that of a typical monomeric αSyn preparation generated by production in *E. coli* and purification under heat-denatured conditions using standard protocols[1-3]. There were no differences in spectra of these two preparations and spectra were typical of those obtained for monomeric αSyn (Fig. S2). To ensure that αSyn remained monomeric in the screen, dynamic light scattering (DLS) analyses were performed on αSyn preparations under screening conditions and in parallel for each library screening experiment. The impact of screening conditions on the state of αSyn was assessed by dynamic light scattering (DLS), a technique highly sensitive for the detection of the formation of higher order oligomers or aggregates. Figure S3 shows volume corrected size distributions of the protein in the screening buffer and properties (mean size/nm, peak area/%) of monomeric peak 1 during a 3 hour incubation, which is the maximum time of αSyn incubation with the microarray during screening. The DLS signals for αSyn were stable over 3 hours under screening assay conditions and generation of oligomers or aggregates was not observed. The area graph of peak 1 reveals that >99.9% of the particle volumes could be attributed to monomeric αSyn at all times investigated. We thus ensured that the HT-CM-SPR screen was performed with monomeric αSyn and thus signals obtained reflected binding of tethered compound to monomeric αSyn.

***Development of αSyn oligomerization assays***

Two novel biochemical assays were developed for αSyn oligomerization based on either bioluminescent complementation of Gaussia luciferase (GLuc) split tags or on FRET of small molecule tags placed onto αSyn. We first established a biochemical assay of αSyn oligomer formation using split GLuc tags as has been used for cellular αSyn oligomerization detection[4]. A mixture of purified αSyn fused at its C-terminus with either the N-Terminal or the C-terminal half of GLuc are incubated and the formation of oligomers monitored by reconstituted GLuc activity

(Figure S5a left panel, blue). For comparison, fibrillization of αSyn was measured in parallel samples by measuring the fluorescence of the dye Thioflavin T (ThT) (Figure S5a left panel black). As expected for an early stage misfolding event, the formation of oligomers did not show the lag phase nor the variability typically seen in the fibrillization assays. The dependence of this assay on αSyn concentrations is shown (Figure S5a right). A FRET based assay for αSyn oligomer formation was developed by tagging αSyn mutagenized to Cys at various locations with Cy3 or Cy5 fluorescent tags and measuring FRET signal from various combinations of tagged proteins using Cy3-tagged αSyn as donor and Cy5-tagged αSyn as acceptor (Figure S5b). Optimal signal was seen with αSyn labeled at position 99 for both donor and receptor molecules. The dependence of this assay on αSyn concentrations is shown in Figure S5c.

### Methods

#### <u>*Protein Expression and purification*</u>

*α-Syn for the HT-CM-SPR screen*. Purified αSyn labeled with $^{15}N$ was purchased from rPeptide (Bogart, GA, USA), and monomeric αSyn further purified by size exclusion chromatography (SEC). The $^{1}H$-$^{15}N$ protein ran as a single peak by SEC, which was collected and stored frozen in single use aliquots for use in all pre-screen analyses and in screening. $^{1}H$-$^{15}N$ HSQC NMR of an aliquot of the purified protein verified that it showed the typical ensemble of αSyn conformations expected for monomeric αSyn[5]. DLS and SDS-PAGE analyses confirmed the monomeric nature of the $^{1}H$-$^{15}N$ αSyn used for the screen.

*α-Syn for the Fibrillization assay.* Wildtype human αSyn was produced from a pET-21d plasmid (Millipore, Burlington, MA) in *E. coli* BL21(DE3) cells induced by addition of 1 mM isopropyl-β-D-1-galactopyranoside and incubated at 16°C overnight. Cells were harvested by centrifugation at 4,000 x g for 10 minutes and stored at -80°C. Frozen cells expressing αSyn protein were resuspended in 50 mM Tris, pH 7.5, 1 mM TCEP, 1 mM EDTA, and 1 mM L-Methionine and lysed using a microfluidizer. Lysate was centrifuged at 7000 x g for 0.5 hours at 21°C. The supernatant was boiled in a water bath for 15 minutes, cooled on ice and centrifuged at 45,000 g for 30 minutes. The supernatant was filtered through a 0.2 µm Corning filter and subjected to ion exchange chromatography using a Capto ImpRes Q (GE Healthcare, Upssala, Sweden) anion exchange (IEX) column at ambient temperature. The column was eluted with a gradient of 100 column volumes from 0 to 1M NaCl using an AKTA Explorer 100 Liquid Chromatography System (GE Healthcare, Uppsala, Sweden). Pure αSyn fractions were desalted to

50 mM $KPO_4$, pH 7.5 and 50 mM NaCl using PD-10 columns (GE Healthcare, Chicago, IL) and concentrated to 5 mg/mL or 10 mg/ml (3K MWCO, Amicon Ultra, Millipore, Burlington, MA).

*α-Syn for the GLuc and FRET oligomerization assays.* For the luciferase complementation oligomerization assay, Gaussia luciferase (GLuc) was split into two non-functional fragments: GLuc1 (residue 18-109) and GLuc2 (residue 110-185). These are the same Gaussia luciferase fragments as the S1 and S2 constructs (originally referred to as syn-hGLuc(1) and syn-hGLuc(2) respectively) used in the cellular αSyn oligomerization assay[6]. Either GLuc1 or GLuc2, without a stop codon, was fused to the C terminus of human αSyn with a flexible linker (IDGGGGSGGGGSSG) placed between αSyn and the GLuc tags. The expression vectors, pET28b-αSyn-GLuc1 (expressing protein αSyn-GLuc1) and pET28b-αSyn-GLuc2 (expressing protein αSyn-GLuc2) were constructed by inserting either coding region into the NcoI/NotI sites in the multiple cloning site (MCS) of pET28b (Novagen). This vector provides His tagged protein as the coding region utilizes the vector stop codon provided after the His tag. For the FRET based oligomerization assay, the human αSyn coding region was amplified using Q5 High-Fidelity DNA Polymerase (NEB) and inserted with its own stop codon into the NcoI/XhoI sites of the pET28b E. Coli expression vector (Novagen). This cloning resulted in no His tag on this protein. Single-point mutations encoding Cys were introduced into αSyn using a Q5 Site-Directed Mutagenesis Kit (NEB). The resultant plasmids (e.g. pET28b-αSyn-Q99C for the position 99 plasmid expressing protein αSyn-Q99C) were used for the recombinant protein expression.

For protein expression for both FRET and GLuc luciferase complementation biochemical assays, BL21(DE3) *E. coli* cells were transformed with the desired plasmids and cultured in LB media at 37°C. At $OD_{600}$ 0.6-0.8, the culture temperature was lowered to 20°C and IPTG (GoldBio) was added to a final concentration of 0.5 mM to induce protein expression and cultures incubated at 20°C overnight. The cells were harvested by centrifugation at 4,000 x g for 20 minutes in an Avanti J-26 XPI centrifuge (Beckman Coulter) with a JLA 8.1000 rotor (Beckman Coulter).

To purify αSyn-GLuc1 and αSyn-GLuc2 proteins, plasmid transformed E coli cells were resuspended in 25 mM Tris, 500 mM NaCl, 0.5 mM TCEP, pH 8.0 and then lysed in EmulsiFlex-C3 (Avestin) in the presence of EDTA-free Protease Inhibitor Cocktail (Roche). The lysate was cleared by centrifugation at 30,000 x g for 30 minutes in a JA 25.50 rotor (Beckman Coulter). His-tagged target protein was purified by Ni-NTA gravity-flow chromatography (Qiagen). Eluted protein was loaded onto MonoQ 10/100 GL (GE) chromatography and eluted with 0-600 mM NaCl gradient. A final purification step was carried out using HiLoad 16/600 Superdex 200 chromatography (GE). The purified protein was filtered through a 0.22 μm filter (E&K Scientific), flash frozen and stored at -80°C.

To purify αSyn used in the FRET assay (αSyn-Q(position number)C; e.g. αSyn-Q99C), plasmid transformed E coli cells were resuspended in 20 mM Tris, pH 8.0 and lysed by boiling for 30 minutes in the presence of EDTA-free Protease Inhibitor Cocktail (Roche). The lysate was cleared by centrifugation at 30,000 x g for 30 minutes in a JA 25.50 rotor (Beckman Coulter). Streptomycin sulfate was added to the lysate at 10 mg/ml to precipitate DNA. After a 30 minutes incubation at 4°C, the lysate was cleared by centrifugation at 30,000 x g for 30 min. Ammonium sulfate was added to the lysate to a final concentration of 0.36 g/ml to precipitate protein. After incubation at 4°C overnight, the protein was pelleted by centrifugation at 30,000 x g for 30 min. The protein was resuspended in 20 mM Tris, 1 mM DTT, pH 8.0, then subjected to MonoQ 10/100 GL (GE) chromatography eluting with 0-600 mM NaCl gradient. A final purification step was carried out using HiLoad 16/600 Superdex 200 chromatography (GE). The monomeric peak was collected and filtered through a 0.22 μm filter (E&K Scientific), flash frozen and stored at -80°C.

*Labeling αSyn with Cy3 and Cy5 dyes*

Purified αSyn-Q(position number)C mutant protein (e.g was reduced by 10 mM DTT for 1 hour at 4°C. The free DTT was removed by HiTrap Desalting chromatography (GE). Cy3-maleimide (GE) or Cy5-maleimide (GE) was added to the reduced αSyn-Q(position number)C at a dye:protein molar ratio of 5:1. The labeling was carried out at 4°C for 12 hours in darkness. The excessive dye was removed by HiTrap Desalting chromatography (GE). The labeled proteins were concentrated using a centricon filter with a 3 kDa cutoff (EMD Millipore). The concentrated protein was filtered through a 0.22 μm filter (E&K Scientific), flash frozen and stored at -80°C.

*Western blotting*

All materials were purchased from Invitrogen (Carlsbad, CA) unless stated otherwise. Cell from a 6 well dish were washed twice with PBS and lysed with 250μl of RIPA Lysis buffer with protease and phosphatase inhibitors (Sigma, St. Louis, MO). Lysates were incubated on a shaker for 15 minutes at 4°C and spun down for 15 minutes at 10,000 x g. Total protein concentrations were determined by BCA assay (Thermo Fisher, Waltham, MA). 1 ml of media was collected and incubated with protease and phosphatase inhibitors and 50 μl of Ni-agarose beads to collect the His-tagged αSyn (Qiagen, Hilden, Germany) for 2 hours at 4°C. Beads were centrifuged and washed in PBS. Cell samples and pelleted beads from the media were placed into Bolt LDS (lithium dodecyl sulfate) sample buffer with 20% β-mercaptoethanol and boiled for 15 minutes. 5 to 15 µg of sample (cells) and an equivalent fraction (of the well) of media were electrophoresed on Bolt 4-12% Bis-Tris Plus gels with Bolt MES SDS running buffer and the Precision Plus Protein Dual Color Standards (Biorad, Hercules, CA) molecular weight markers. After electrophoresis, the separated proteins were transferred onto a 0.2 µm pore size PVDF membrane for 90 minutes at 400 mA at 4°C in Bolt Transfer Buffer using BioRad midi transfer chambers. Post-transfer, membranes were treated for 30 minutes with 0.4% PFA in PBS to enhance synuclein binding[7] rinsed with water, stained with 0.1% Ponceau S in 5% acetic acid, rinsed with Phosphate Buffered Saline with Tween 20 (PBST), and blocked in Odyssey PBS Blocking Buffer (LI-COR, Lincoln, NE) for 1 hour at room temperature. Membranes were then incubated with primary antibody in Odyssey PBS Blocking Buffer with 0.1% Tween 20 overnight at 4°C. Purified mouse 5c12 antibody[8] detecting total αSyn was diluted 1/1000, and mouse anti actin antibody clone AC15, (Sigma-Aldrich, St. Louis, MO) was diluted at 1/35,000. Membranes were then washed 4 times for 10 minutes in PBST and incubated with secondary antibody goat anti-mouse infra red 800 (LI-COR, Lincoln, NE) diluted at 1/10 000 in Odyssey PBS Blocking Buffer with 0.2% Tween 20 followed by washing 4 times for 10 minutes in PBST. Membranes were scanned and quantitated using the Odyssey CLx Imaging System (LI-COR, Lincoln, NE). Actin and αSyn were visualized in the 800 nm fluorescent channel and the molecular weight markers visualized at 700 and images merged. In cases where the image is cut to remove irrelevant lanes, a bar is placed in the image.

*DLS Analyses of αSyn*

To confirm its monomeric state and to check the integrity of the protein, DLS measurements were performed with a Malvern Zetasizer Nano instrument. In order to remove high molecular weight impurities all buffers used in the DLS studies were filtered through a 20nm filter and the concentrated αSyn stock, at 1.2 mg/ml in PBS, was filtered through a 100kDa cut-off spin device. Prior to screening on library arrays the protein stock solution was freshly diluted down into the selected screening buffer to a concentration of 9 µM and amounts of monomeric/oligomeric species were monitored over the three hour course of the array experiments. From autocorrelation plots intensity and volume corrected plots were calculated.

*SPR Screening of Monomeric αSyn*

*Chemical Microarrays:* The construction of the arrays and their use for primary screening in drug discovery was described elsewhere[9,10]. All 114,000 library compounds, the synthesis and QC of which was described before[11], were coupled to a flexible, long, hydrophilic thiol-linker[11]. Upon pintool spotting, the linker-compound constructs were allowed to react covalently with maleimide moieties present in a mixed self-assembled monolayer (SAM) surface on the array surface. Eventually, a surface architecture consisting of glass/gold/SAM surface/covalently attached chemtag/immobilized ligands was achieved. The ligand density was adjusted by varying the ratio of maleimide-attached (anchor) thiols to unmodified (diluent) thiols in the mixed SAM. The optimized surface chemistry was designed to be resistant to nonspecific protein binding exhibiting only marginal background in the SPR screening experiments. Each microarray contained 9,216 sensor fields corresponding to different tethered sample spots on the array.

*SPR Imaging of Chemical Microarrays:* Intermolecular interactions during the HT-CM-SPR were detected by recording the shift in the wavelength dependent surface plasmon resonance (SPR) minima of the chemical microarrays upon αSyn binding. The microarrays were analysed using *NovAliX*'s (previously Graffinity Pharmaceuticals's) in-house developed SPR Imager® instrument. The optical set-up in the instrument allowed illuminating the entire chip area with parallel light of defined incidence angle and wavelength via a high refractive index prism in a Kretschmann configuration. Reflection images of the chip were recorded by means of a cooled, low-noise CCD camera. While keeping the incidence angle of the incoming beam fixed, the wavelength was varied over a range covering the SPR resonance conditions for the given chip/prism combination. Recorded array images were deconvoluted by automatic spot finding routines and grey scale analysis. Plotting the reflectivity of the individual sensor areas versus the applied wavelength yielded 9,216 SPR minima for each microarray resulting from the excitation of Surface Plasmons associated with sample spots on the arrays. Binding of analytes to the immobilized library compounds altered the optical resonance conditions for the corresponding sensor fields. Differences from that of analyte free buffer were detected by monitoring (red) shifts of the wavelength dependent SPR minima with time. Additionally, bulk refractive index changes upon buffer exchange were taken into account by control spots distributed across the array. Typical incubation times of analytes on the arrays ranged from 15 minutes to 3 hours during which scans were recorded repeatedly. SPR signals were visualized in coloured 2D fingerprints for manual hit selection using JARRAY, NovAliX's proprietary software for visualizing chemical microarray data. Manual hit picking of individual library arrays was completed by detailed data mining performed across all screened arrays.

*Hit selection and compound resyntheses*: A software routine guided the hit selection process on the array level by fitting Gaussian functions to the SPR signal distribution and suggesting hit thresholds per array allowing removal of non-hits. A great deal is known about the interaction of library components tethered on the chip with other proteins from prior screens[9,11-14]. Therefore, compounds showing promiscuous interactions (frequent binders identified to have hit > 50% of screened targets) were excluded from initial hits to extract compounds with possibly higher target specificity.

The 65 resynthesized compounds chosen for testing in functional assays were verified as the indicated structure and of sufficient purity by $^1$H-NMR and by liquid chromatography and mass spectrometry analyses (LC-MS). $^1$H NMR analyses were performed on an AVANCE III 400 HD, 400 MHz with NS = 4, DMSO-$d_6$ as solvent. Spectra analyses were performed using either TOPSPIN or MNova software. These 65 compounds were at least 85% pure, with most over 95% pure. The active compounds were all over 90% pure. The LC-MS determined purity and $^1$H-NMR peaks for the 9 active compounds described herein are below:

573416

$^1$H NMR (400 MHz, DMSO-$d_6$) δ 8.16 (d, J=8.11 Hz, 1 H),7.93-7.91(m, 2H), 7.79 (d, J=8.15 Hz, 1 H),7.57-7.49(m, 2H),7.45-7.42(m, 1H),7.35-7.33(m, 1H),3.39-3.32(m, 1H),3.14(dd,J=13.32,5.96Hz,1H)2.93(dd,J=13.34,7.46Hz,1H), 2.56 (d, J=4.61 Hz, 3H),2.22-2.11(m, 2H),1.84(s,2H); LCMS(m/z): [M]+ calcd. for C15 H18 N2 O, 242.32; found, 243.2; abundance 97.92%.

573417

$^1$H NMR (400 MHz, DMSO-$d_6$) δ 9.33 – 9.25 (m, 6H), 8.74 (d, *J* = 4.4 Hz, 1H), 8.62 (d, *J* = 5.5 Hz, 3H), 8.52 (d, *J* = 8.5 Hz, 1H), 8.15 – 8.06 (m, 6H), 7.94 (s, 9H), 7.81 (dddd, *J* = 33.4, 8.4, 6.9, 1.4 Hz, 7H), 7.69 (s, 3H), 7.52 (s, 3H), 7.49 (d, *J* = 4.6 Hz, 0H), 6.57 (s, 1H), 4.81 (td, *J* = 8.3, 4.9 Hz, 3H), 3.42 – 3.34 (m, 2H), 3.22 (dd, *J* = 13.0, 8.1 Hz, 3H), 2.55 (s, 15H); LCMS(m/z): [M]+ calcd. for C13 H14 N4 O2, 258.28; found, 259.00; abundance >99%.

573418

$^1$H NMR (400 MHz, DMSO-$d_6$) δ 8.86 (d, *J* = 8.3 Hz, 1H), 8.66 (dt, *J* = 4.7, 1.3 Hz, 1H), 8.59 (s, 1H), 8.05 – 7.95 (m, 2H), 7.67 – 7.58 (m, 2H), 7.31 (s, 1H), 7.19 (d, *J* = 1.3 Hz, 1H), 4.73 (td, *J* = 8.0, 5.0 Hz, 1H), 3.17 (qd, *J* = 15.1, 6.5 Hz, 2H), 2.54 (s, 1H); LCMS(m/z): [M]+ calcd. for C12 H13 N5 O2, 259.268; found, 260.2; abundance 98.710%.

573419

$^1$H NMR (400 MHz, DMSO-$d_6$) δ 8.36 (s,1 H), 8.03 (m, 3 H),7.69(s,1 H),7.59-7.53(m, 3 H),7.26(s,1 H), 4.39(dd, J=9.14,6.66Hz,1H),1.80-1.59(m, 6H),1.20-0.96(m, 6H); LCMS(m/z): [M]+ calcd. for C18 H21 N3 O2 S, 343.449; found, 344.05; abundance 99.810%.

573434

$^1$H NMR (400 MHz, DMSO-$d_6$) δ 10.07(s,1H),9.77(s,1H), 7.61 (dd, J = 7.64, 1.10 Hz, 1H), 7.46(s,1H),7.41(m, 1H), 7.32 (td, J = 7.44, 1.18 Hz,1H),4.40(dd, J

=3.62,13.34Hz,1H),4.12(dd, J =7.62,13.34Hz,1H),3.21-3.16(m,1H), 2.58 (d, J=5.01 Hz, 3H),2.11-
2.07(m,5H), 1.84 (t, J=11.22 Hz, 2H), 1.62 (d, J=12.38 Hz, 1 H),1.55-1.48(m,2H),1.27-1.10(m,3H); LCMS(m/z): [M]+ calcd. for C16 H24 N2 O, 260.379; found, 261.1; abundance 93.986%.

573437

$^1$H NMR (400 MHz, DMSO-$d_6$) δ 8.40 (s, 1H), 8.02 (d, *J* = 8.4 Hz, 1H), 7.91 – 7.84 (m, 1H), 7.79 (d, *J* = 8.0 Hz, 1H), 7.65 (s, 1H), 7.44 – 7.30 (m, 2H), 7.23 (s, 1H), 6.92 (s, 1H), 4.25 (t, *J* = 7.5 Hz, 1H), 3.20 (t, *J* = 12.5 Hz, 2H), 2.73 (q, *J* = 11.7 Hz, 2H), 2.55 (d, *J* = 10.2 Hz, 4H), 1.98 (s, 1H), 1.83 – 1.70 (m, 2H), 1.53 (q, *J* = 12.6 Hz, 2H); LCMS(m/z): [M]+ calcd. for C19 H22 N4 O S2, 386.542; found, 387.05; abundance 91.350%.

576755

$^1$H NMR (400 MHz, DMSO-$d_6$) δ 8.54 (s, 1H), 8.06 (d, *J* = 4.7 Hz, 1H), 6.83 (d, *J* = 2.8 Hz, 1H), 6.65 (dd, *J* = 8.7, 2.7 Hz, 1H), 6.53 (d, *J* = 8.6 Hz, 1H), 5.69 (s, 2H), 2.68 (d, *J* = 4.5 Hz, 3H), 2.53 (d, *J* = 0.5 Hz, 1H); LCMS(m/z): [M]+ calcd. for C8 H10 N2 O2, 166.179; found, 167.15; abundance 98.43%.

581674

$^1$H NMR (400 MHz, DMSO-$d_6$) δ 8.18 (d, *J* = 3.0 Hz, 1H), 8.02 (d, *J* = 5.4 Hz, 2H), 7.74 – 7.64 (m, 8H), 7.56 – 7.42 (m, 9H), 7.42 – 7.33 (m, 2H), 4.61 (t, *J* = 7.0 Hz, 2H), 3.17 (d, *J* = 0.6 Hz, 1H), 2.74 (dd, *J* = 6.9, 4.0 Hz, 4H), 2.55 (t, *J* = 4.5 Hz, 6H); LCMS(m/z): [M]+ calcd. For C16 H18 N2 O, 254.331; found, 255.1; abundance 99.82%.

582032

$^1$H NMR (400 MHz, DMSO-$d_6$) δ 7.91 (d, *J* = 4.8 Hz, 1H), 7.84 (dt, *J* = 8.0, 1.0 Hz, 1H), 7.72 – 7.68 (m, 1H), 7.63 (overlap, 1H), 7.43 – 7.36 (m, 4H), 7.33 – 7.29 (m, 1H), 5.75 (q, *J* = 7.1 Hz, 1H), 4.28 (d, *J* = 16.6 Hz, 1H), 4.20 – 4.10 (overlap, 2H), 3.76 (d, *J* = 17.9 Hz, 1H), 2.60 (d, *J* = 4.6 Hz, 3H), 1.39 (d, *J* = 7.1 Hz, 3H); LCMS(m/z): [M]+ calcd. for C20 H20 Cl N3 O3, 385.849; found, 386; abundance 95.00%.

The following software was used to calculate parameters of hit compounds: Hivol for the calculation of the counts, Biobyte for the calculation of ClogP and Openbabel v. 2.3.1 for the calculation of TPSA.

*Statistical Analysis of compound activity in Fibrillization assays.*

For the analyses of compound activity in the αSyn fibrillization assay, the relative fluorescence unit (RFU) data obtained from compounds were compared to those of DMSO, to identify those that are statistically different from DMSO at various time points. Each experiment contained 4 replicates of each compound and of DMSO control. To identify compounds with significantly different RFU from DMSO, results at hours 0, 3, 50, 60, 74, 82, and 112 were examined. A repeated measures analysis of variance was performed on logarithmic transformed data using a linear mixed effects model which included Compound, Time, Replicate (Compound) and

Compound*Time. The covariance structure across time that minimized the corrected Akaike Information Criterion (AICC)[15] was used and the Kenward and Roger method[16] was used to determine the appropriate degrees of freedom. Each compound mean was compared to the DMSO mean using Bonferroni multiplicity adjusted t-tests. Statistical analyses were conducted using the MIXED procedure in SAS 9.1 (SAS Institute Inc. Cary, NC, USA.,[17]). Statistical significance refers to p-value < 0.05.

*Dopaminergic Neuron Viability Assay*

Primary midbrain cultures were prepared and dopaminergic neuronal viability in the presence and absence of αSyn was assessed as previously described[18,19] and repeated herein. Primary midbrain cultures were prepared from embryonic day 17 embryos of Sprague-Dawley rats using methods reviewed and approved by the Purdue Animal Care and Use Committee as described previously[18,19]. After dissection, the cells were plated into a poly-L-lysine-treated 48-well plate at a density of 163,500 cells per well. Four days later, the cells were treated with cytosine arabinofuranoside (AraC) (20 µM) for 48 hours to inhibit the growth of glial cells. The cultures were used for measurements of dopaminergic neuron viability at 7 days in vitro. Primary midbrain cultures were transduced with αSyn A53T adenovirus (MOI = 10), in the absence or presence of ELN576755, as described previously[18,19]. After 72 hours, the cells were incubated with fresh media with or without the compound for another 24 hours prior to immunocytochemical analysis, which was carried out as described previously[18,19]. The cells were fixed, permeabilized, and blocked prior to an overnight treatment with two primary antibodies: a mouse monoclonal IgG specific for microtubule-associated protein 2 (MAP2) (1:500) and a rabbit polyclonal antibody specific for tyrosine hydroxylase (TH) (1:500). Next, the cells were treated with two secondary antibodies, goat anti-mouse IgG conjugated to AlexaFluor 594 (1:1000) and goat anti- rabbit IgG conjugated to AlexaFluor 488 (1:1000) for 1 hour. In order to determine the viability of dopaminergic neurons, MAP2- and TH-positive neurons were counted in 10 to 15 randomly chosen observation fields in a blinded manner using a Nikon TE2000-U inverted fluorescence microscope (Nikon Instruments, Melville NY) with a 20X objective. In the control conditions, and in conditions where the compound was neuroprotective, we typically counted 500 to 1,300 MAP2-positive neurons, a range that corresponds to 20 to 50 TH–positive neurons[18,20]. The data were expressed as the ratio of the TH-positive neurons to the MAP2-positive neurons. Each experiment was repeated at least three times using embryonic midbrain cultures from different pregnant rats.

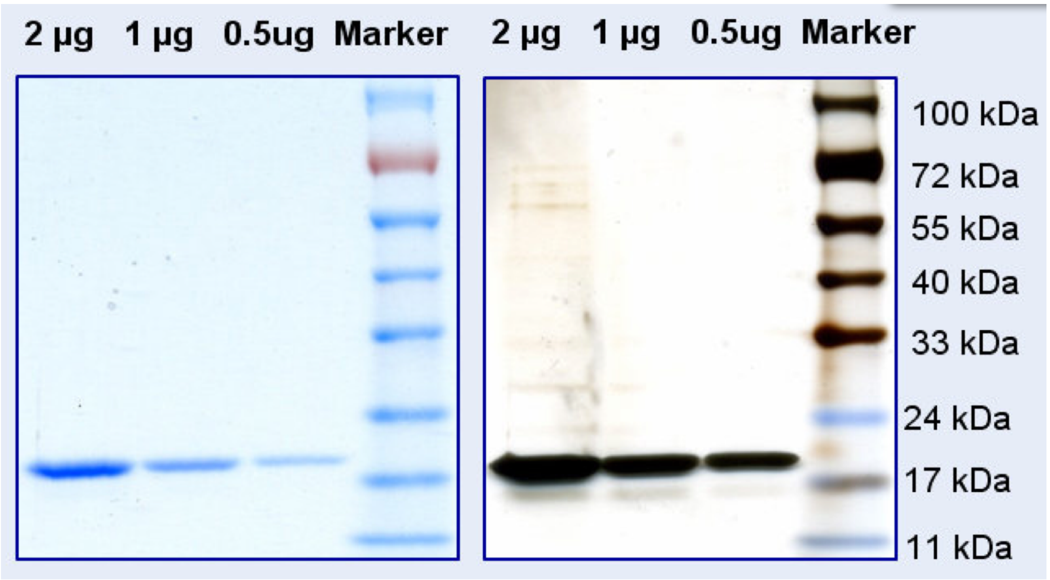


**Figure S1. SDS-PAGE analyses of αSyn used for HT-CM-SPR screen.** Sodium dodecyl sulfate polyacrylamide gel electrophoresis (SDS-PAGE) followed by staining of proteins with Coomassie (left) or silver (right) was performed to evaluate purity and size of αSyn. αSyn is pure and predominantly monomeric in size as detected in this electrophoresis system. Mass Spectrometric analyses of this preparation showed it to be of > 99% purity and without detectable degradation products (data not shown).

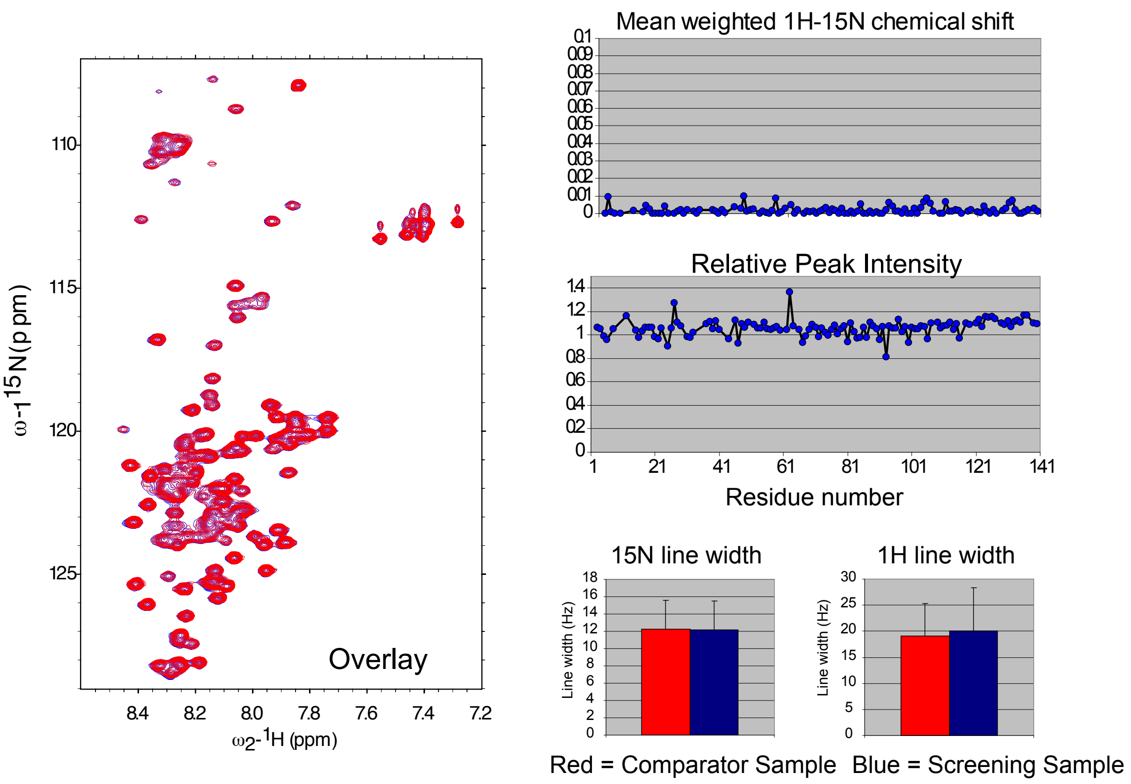


**Figure S2. H[1]-N[15]-HSQC analyses of αSyn used for screen and comparator monomeric αSyn preparations.** NMR spectroscopic techniques were as described[2]. Overlaying the spectra of the screening sample (blue) and comparator (red) preparations show identical spectra (left). The chemical shifts and relative peak intensities of each amino acid and $^{15}$N and $^{1}$H line widths (right) are identical.

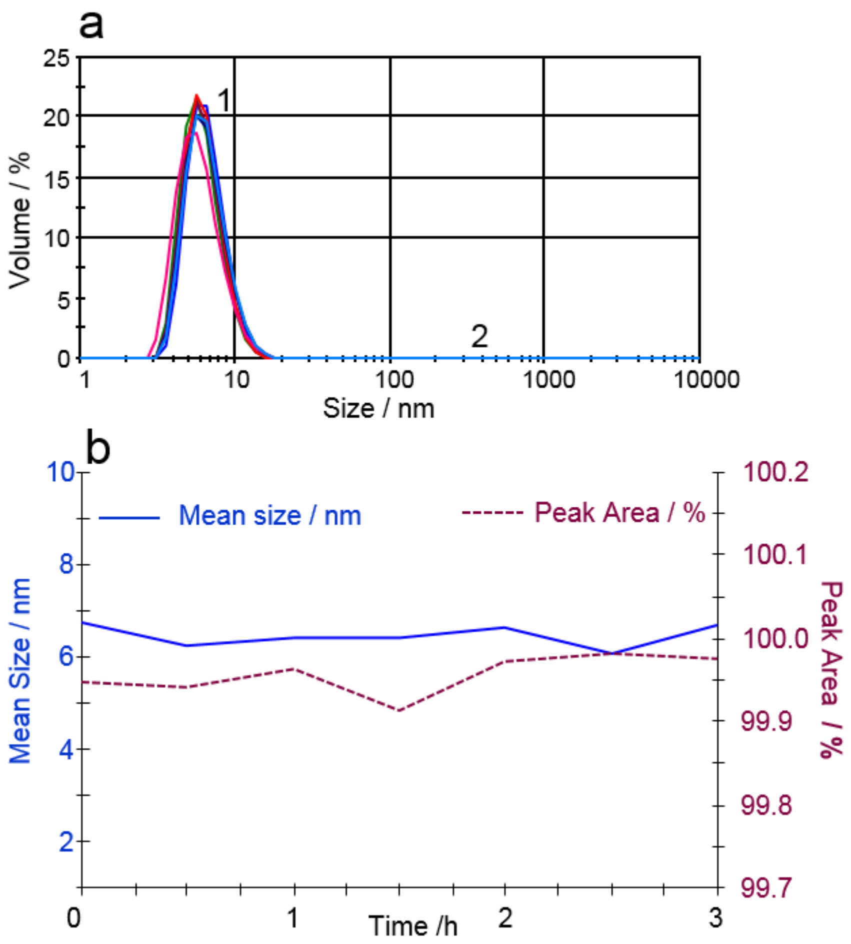


**Figure S3. DLS measurements of αSyn used for the HT-CM-SPR screen indicate monomeric structural state composition.** DLS analyses of the αSyn sample used in the HT-CM-SPR screen were done after incubation for up to 3 hours under screening conditions to explore its heterogeneity. **a)** Volume corrected size distributions of the αSyn sample are shown with measurements at different incubation times overlaid. The αSyn sample was principally monomeric αSyn (peak 1), while negligible amounts of oligomeric αSyn (peak 2) were observed. **b)** The size (hydrodynamic diameter) and area percentage (Area %) of Peak 1, corresponding to monomeric αSyn, were stable for over 3 hours, and >99.9% of the particle volume could be attributed to monomeric αSyn in the sample.

576755

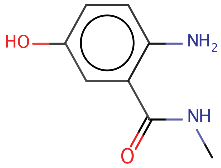

582032

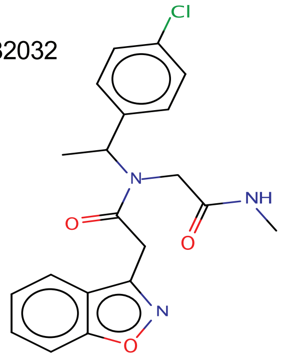

573437

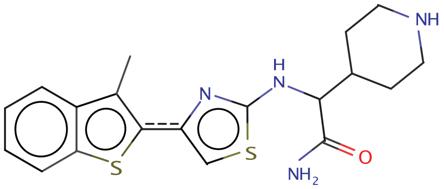

573416

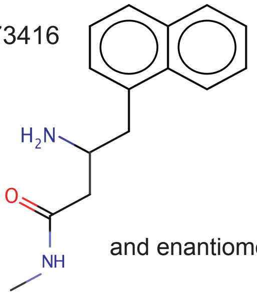

and enantiomer

573417

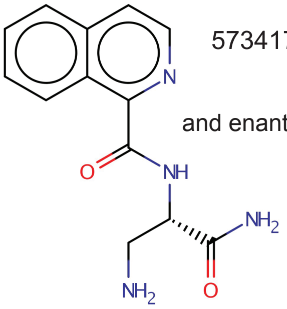

and enantiomer

573418

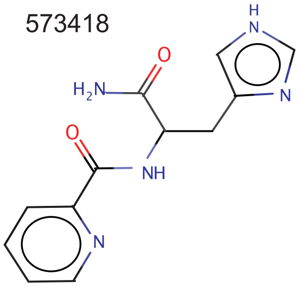

573419

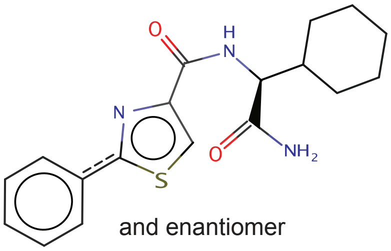

and enantiomer

581674

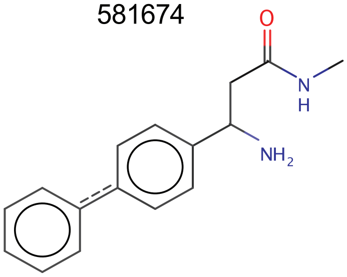

573434

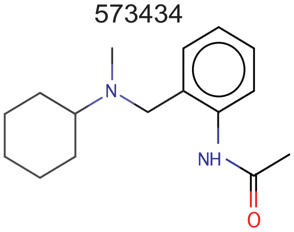

**Figure S4. Structures of active compounds.**

IUPAC names for the shown compounds are below:

573416: (R)-3-Amino-N-methyl-4-naphthalen-1-yl-butyramide
573417: Isoquinoline-1-carboxylic acid ((S)-2-amino-1-carbamoyl-ethyl)-amide
573434: N-{2-[(Cyclohexyl-methyl-amino)-methyl]-phenyl}-acetamide
573437: 2-[4-(3-Methyl-benzo[b]thiophen-2-yl)-thiazol-2-ylamino]-2-piperidin-4-yl-acetamide
573418: Pyridine-2-carboxylic acid [(S)-1-carbamoyl-2-(1H-imidazol-4-yl)-ethyl]-amide
573419: 2-Phenyl-thiazole-4-carboxylic acid ((S)-carbamoyl-cyclohexyl-methyl)-amide
576755: 2-Amino-5-hydroxy-N-methyl-benzamide
581674: 3-Amino-3-biphenyl-4-yl-N-methyl-propionamide
582032: 2-Benzo[d]isoxazol-3-yl-N-[1-(4-chloro-phenyl)-ethyl]-N methylcarbamoylmethyl-acetamide

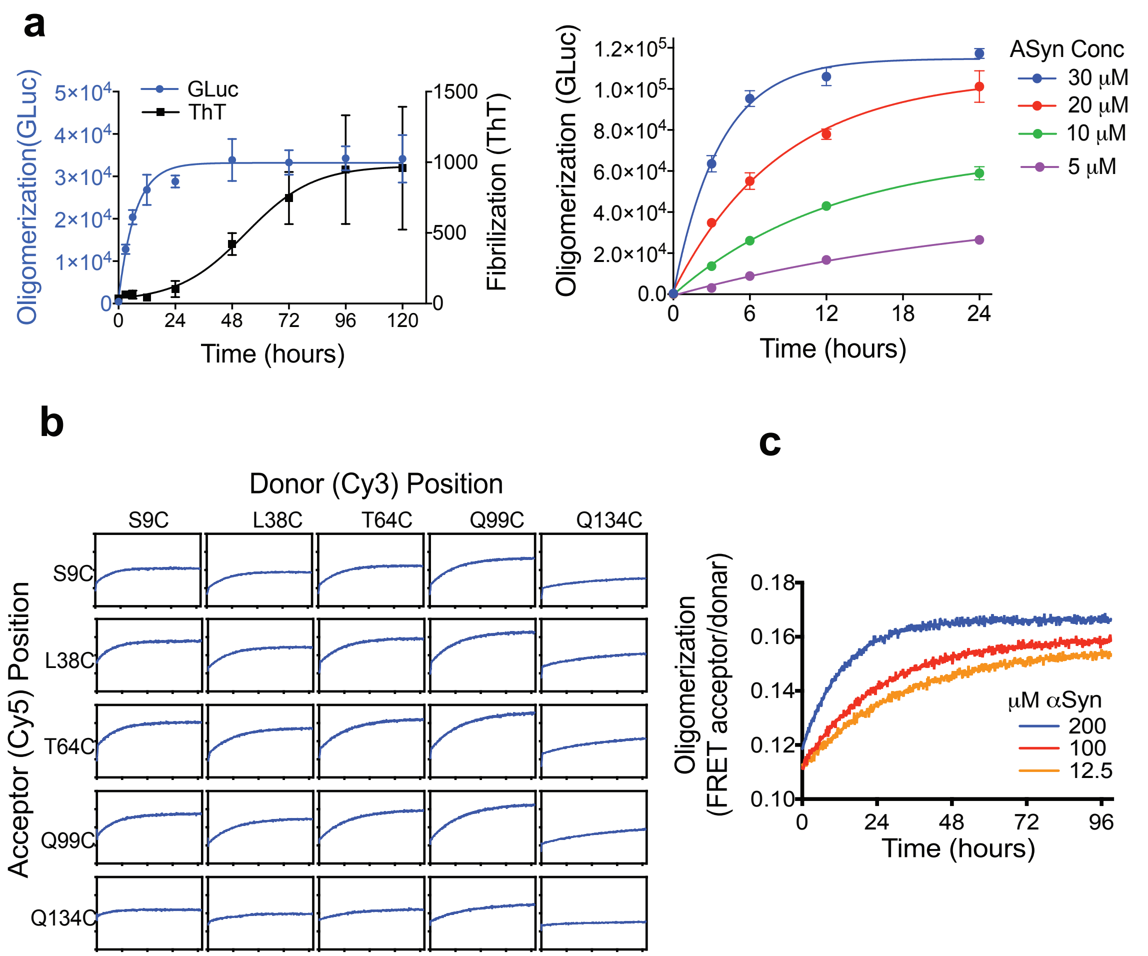


**Fig S5. Development of biochemical αSyn oligomerization assays.**

**a)** Left: The bioluminescent complementation assay (blue) is shown compared to a thioflavin T (ThT) fluorescent based fibrillar assay (black). N and C terminal split Gluc tagged onto the C-Terminus of αSyn were incubated with shaking and aliquots assayed at various times for oligomerization as described in the Methods. Parallel samples were assayed for fibrillization by addition of 25 µM Thioflavin T and reading using excitation at 440 nm and emission 482 nm. Mean +/- SD shown. Right: Varying concentrations of split Gluc tagged αSyn were incubated and assayed for oligomerization. For this panel incubation was performed in a PCR machine without shaking at 37$^{o}$C. **b)** Development of the FRET αSyn oligomerization assay. αSyn mutagenized to Cys at the indicated location was labeled with Cy3 or Cy5 dyes as described herein and oligomerization measured as described in Methods of the associated paper. Optimal FRET signal was seen with Q99C location as both donor and receptor. **c)** The dependence of the FRET oligomerization assay on αSyn concentrations is shown.

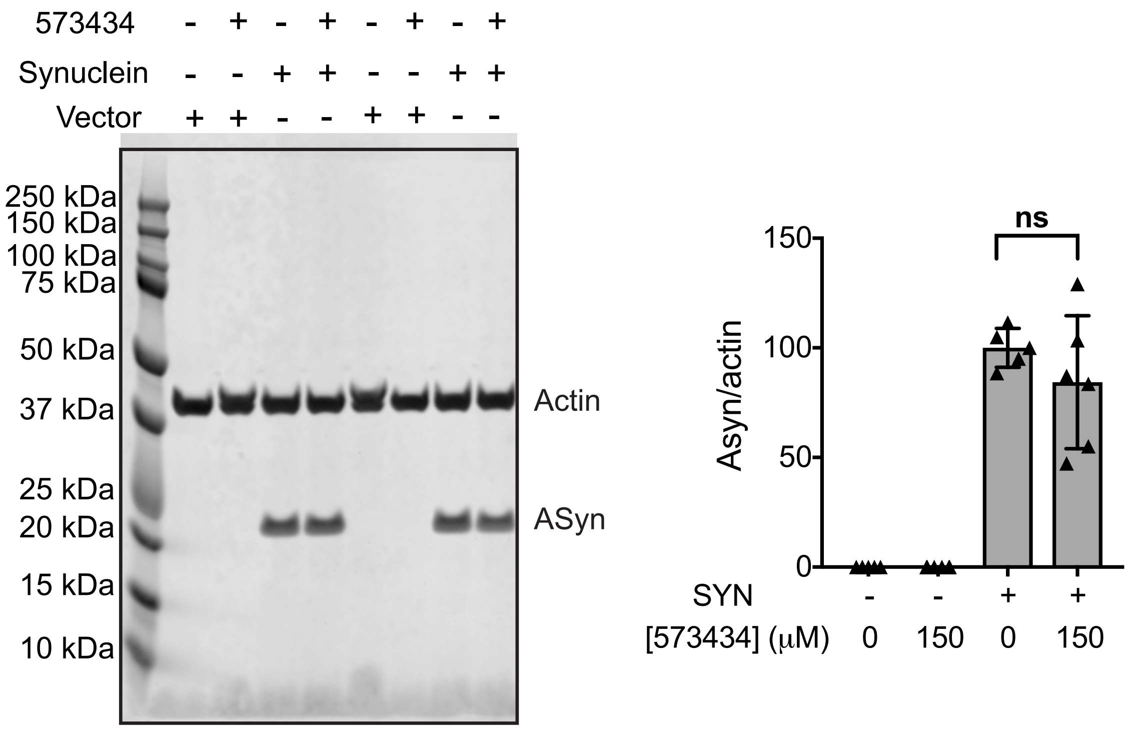


**Fig S6. Western analyses of αSyn levels in B103 cells treated with 573434.** Left: Representative Western blot showing αSyn levels in cells of one experiment with duplicate wells for each condition. This is a separate experiment from the Western image shown in Fig. 5e of the original manuscript. Entire length of blot shown. Outline of full blot shown by black lines. Right: Westerns from multiple experiments were quantitated and the αSyn band intensity from cells was normalized to that of actin. Three experiments with duplicate wells are combined in this figure. Each data point is a separate well. These are the same experiments for which media/cell values are calculated in Fig. 5e. There were no statistical differences between control and 573434 treatment by unpaired t test (alpha=.05). Mean ± SD.

| Molecular Weight Distribution | Number of hits |
|---|---|
| fragments < 250 Da | 13 |
| 250 Da < fragments < 300 Da | 31 |
| 300 Da < lead-like < 350 Da | 42 |
| 300 Da < lead-like < 400 Da | 39 |
| 400 Da < lead-like< 500 Da | 27 |

**Table S1. Molecular weight distribution of 152 selected hits identified by the HT-CM-SPR of monomeric αSyn.** Molecular weight calculated as in table 1.
